# A *trans-*acting long non-coding RNA represses flowering in *Arabidopsis*

**DOI:** 10.1101/2021.11.15.468639

**Authors:** Yu Jin, Maxim Ivanov, Anna Nelson Dittrich, Andrew D. L. Nelson, Sebastian Marquardt

**Affiliations:** Department of Plant and Environmental Sciences, Copenhagen Plant Science Centre, University of Copenhagen, Frederiksberg, Denmark; Boyce Thompson Institute, Cornell University, Ithaca, NY, USA

**Keywords:** lncRNA, *FLAIL*, *trans-*acting, chromatin binding, flowering regulation, ChIRP-seq

## Abstract

Eukaryotic genomes give rise to thousands of long non-coding RNAs (lncRNAs), yet the purpose of lncRNAs remains largely enigmatic. Functional characterization of lncRNAs is challenging due to multiple orthogonal hypothesis for molecular activities of lncRNA loci. Here, we identified a **fl**owering **a**ssociated **i**ntergenic **l**ncRNA (*FLAIL*) that represses flowering in *Arabidopsis*. An allelic series of *flail* loss-of-function mutants generated by CRISPR/Cas9 and T-DNA mutagenesis showed an early flowering phenotype. Gene expression analyses in *flail* mutants revealed differentially expressed genes linked to the regulation of flowering. A genomic rescue fragment of *FLAIL* introduced in *flail* mutants complemented gene expression defects and early flowering, consistent with *trans-*acting effects of the *FLAIL* RNA. Knock-down of *FLAIL* RNA levels using the artificial microRNA approach revealed an early flowering phenotype shared with genomic mutations, indicating a *trans-*acting role of *FLAIL* RNA in the repression of flowering time. Genome-wide detection of *FLAIL*-DNA interactions by ChIRP-seq suggested that *FLAIL* may directly bind genomic regions. *FLAIL* bound to genes involved in regulation of flowering that were differentially expressed in *flail*, consistent with the interpretation of *FLAIL* as a *trans-*acting lncRNA directly shaping gene expression. Our findings highlight *FLAIL* as a *trans-*acting lncRNA that affects flowering in *Arabidopsis*, likely through mediating transcriptional regulation of genes directly bound by *FLAIL*.

## Background

The purpose of DNA sequences that do not encode proteins represents an open question in the biology of genomes. It is now clear that RNA polymerase II (RNAPII) converts non-coding DNA into non-coding RNA genome-wide [1]. Long non-coding RNAs (lncRNAs) are key products of ubiquitous RNAPII transcription in the non-coding genome [2]. Non-coding DNA engaged in lncRNA production may function through various molecular mechanisms, ranging from roles as DNA elements, RNAs, the act of transcription and small peptides [3, 4]. This wide range of possible cellular roles affects strategies to elucidate their functions experimentally [5]. Functional characterization of DNA elements benefits from a high-quality annotation of lncRNAs in genomes [6]. Integration of several orthogonal transcriptomic data offers an opportunity to inform on the precise location of various lncRNA subtypes and alternative lncRNA isoforms from a single locus [6]. Experimental avenues to abolish specific lncRNA isoforms may trigger the generation of alternative isoforms that may partially substitute for functions, calling for a multi-facetted functional characterization of lncRNA loci [7, 8]. The resulting RNA molecules, but also the act of transcription generating lncRNA can regulate the expression of neighboring genes [9]. The act of transcription from an upstream lncRNA locus may trigger gene activation of the downstream, or gene repression, for instance by transcriptional interference [9]. Ribosome profiling (Ribo-seq) may suggest small open reading frames (sORFs) of the lncRNA loci [3]. Notably, lncRNA association with ribosomes may indicate either ribosome-coupled RNA degradation or translation [3]. Collectively, the broad range of candidate hypotheses by which lncRNA loci may play functional roles call for multiple approaches to distinguish alternative molecular mechanisms [10].

LncRNAs may regulate nearby genes in *cis* or distant genes in *trans* [11–13]. Compared to *cis-*acting lncRNAs, relatively fewer functions of *trans-*acting lncRNAs have been clarified [4]. A key experiment to distinguish between *cis-*acting and *trans-*acting mechanisms is to test phenotypic complementation of lncRNA loss-of-function mutants by lncRNA expression from a different genomic region [4]. *Trans-*acting lncRNAs regulate distant genes via different mechanisms [11], for example, through chromatin targeting of lncRNAs to fine tune chromatin architecture resulting in an altered transcriptional output [14]. Nevertheless, functional characterization of *trans-*acting lncRNAs remains a key knowledge gap to understand the regulatory contributions of the non-coding genome.

In plants, an increasing number of lncRNA loci have been implicated in the regulation of flowering time [15–20]. Flowering time represents a developmental transition of plants that is key for reproductive success. Genetic and environmental factors, for example, altered internal secondary metabolites (e.g. lignin), extended cold periods (i.e. vernalization) or day length (i.e. photoperiod), help plants to align flowering with favorable conditions [21–23]. Vernalization-induced flowering associates with several lncRNAs such as *COOLAIR*, *COLDAIR*, *ANTISENSE LONG* (*ASL*), and *COLDWRAP* that in *cis* repress gene expression of *FLOWERING LOCUS C* (*FLC*), a key flowering repressor at different stages of vernalization [24–27]. The contribution of *trans-*acting lncRNAs to the regulation of flowering time is currently unclear.

Here, we characterized the lncRNA locus *FLAIL* in *Arabidopsis* that gives rise to several RNA isoforms. Genomic mutations and strand-specific RNA repression provided evidence that *FLAIL* sense lncRNA repressed flowering. Genetic complementation data supported a *trans-*acting role of *FLAIL* in the regulation of flowering genes. *FLAIL* RNA bound the chromatin of flowering-related target genes that were differentially expressed in *flail* mutants, arguing for direct effects of *FLAIL* in flowering gene regulation. In summary, our data suggest flowering regulation through effects on gene expression by chromatin association of the *trans-*acting RNA *FLAIL*.

## Results

### Characterization of the *FLAIL* locus

*FLAIL* was annotated as a lncRNA in the TAIR [28] and GreeNC [29] databases and mapped as a single exon to chromosome 2 in *Arabidopsis* (Fig. 1A). Consistently, Nanopore RNA-seq data of chromatin-associated RNAs provided no evidence for splicing at the *FLAIL* locus [30] (Fig. S1A). Additionally, plaNET-seq used for genome-wide profiling of nascent RNA polymerase II (RNAPII) transcription [31], identified both sense and antisense isoforms at the *FLAIL* locus (Fig. 1A). We obtained additional information by examining RNA 5′-end mapping by TSS-seq [32] and simultaneous RNA 5′- and 3′-end mapping by TIF-seq [33] in the mutant of the nuclear exosome component *HUA ENHANCER2* (i.e. *hen2-2*). These data confirmed that the *FLAIL* locus was transcribed on both strands, since sense and antisense *FLAIL* transcripts were detected. Moreover, RNA isoforms derived from *FLAIL* were 5′-end capped, 3′-end polyadenylated and degraded by the nuclear exosome in *Arabidopsis* (Fig. 1A). To assess the protein-coding potential of *FLAIL* RNAs, we used the Coding Potential Calculator (CPC2) [34] and Coding-NonCoding Identifying Tool (CNIT) [35]. These analyses revealed poor coding potential of the lncRNA *FLAIL*, similar to other well-known ncRNAs (*18sRNA*, *U6*, *ELENA1*, *SVALKA*) [36, 37], but much lower than the protein coding potential of *UBIQUITIN* (*UBQ*) mRNA (Fig. 1B). Finally, an analysis of translation start sites in *FLAIL* RNAs suggested poor protein coding potential, well below the 0.5 threshold predicted for protein start codon as determined by the NetStart software package [38] (Fig. S2). Nevertheless, Ribo-seq [39, 40] data indicated ribosome association of *FLAIL* RNAs. This ribosome association could be consistent with translation of two sORFs with ∼9 amino acids, or ribosome-mediated RNA degradation (Fig. S1B) [39]. In summary, the *FLAIL* locus harbors both sense and antisense lncRNA isoforms.

**Fig. 1.**
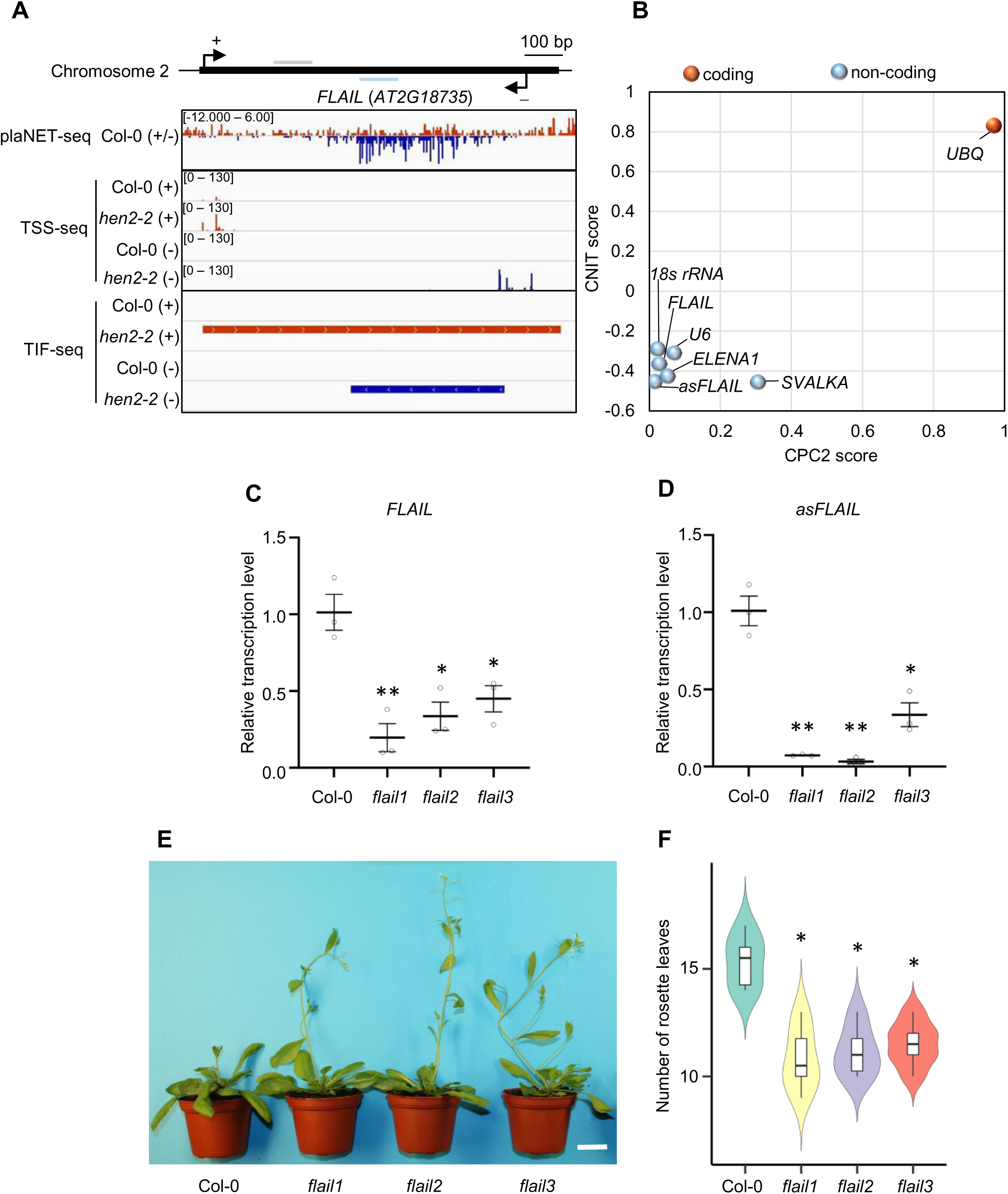
Phenotypes of *FLAIL* knock-down plants. **A** Genome browser screenshot of plaNET-seq, TSS-seq and TIF-seq at the *FLAIL* genomic region in Col-0 and *hen2-2* mutant. Sense (+) and antisense (-) strands were shown in red and dark blue, respectively. Grey bar and light blue bar indicated primer locations of RT-qPCR for sense *FLAIL* (*FLAIL*) and antisense *FLAIL* (*asFLAIL*), respectively. **B** Coding potential of the transcript in the genomic region of *FLAIL* and *asFLAIL* and reference transcripts including non-coding RNAs (*18sRNA*, *U6*, *ELENA1*, *SVALKA*) and coding gene *UBQ* according to the CNIT and CPC2 algorithm. **C-D** Detection of *FLAIL* and *asFLAIL* gene expression in Col-0 and *flail* mutants by RT-qPCR. Transcript levels were normalized to *UBQ* expression levels. Y-axis showed relative values compared to the expression level of Col-0. Bars represented average ± s.e.m (n = 3 independent 14-d seedling pools). *, *p* value < 0.05 and **, *p* value < 0.01 by Student’s t-test compared to Col-0. **E** Morphological phenotypes of 4-week-old plants of Col-0, *flail* mutants at 20 °C under a 16-h light/8-h dark growth condition. Scale bar: 2 cm. **F** Violin graph showed number of rosette leaves after appearance of the first flower bud in Col-0. Data represented the mean of six independent experiments. Boxes spanned the first to third quartile, bold black lines indicated median value for each group and whiskers represented the minimum and maximum values. *, *p* value < 0.05 was indicated by Student’s t-test compared to Col-0.

Even though lncRNAs show relatively poor sequence conservation [41], *trans*-acting lncRNAs may show signatures of conservation across species [42]. We identified a match in the genome of *Camelina sativa* on chromosome 17 (Fig. S3A). *Camelina*, like *Arabidopsis*, is in the Brassicaceae and last shared a common ancestor with *Arabidopsis* ∼ 18 million years ago [43]. Reciprocal blast searches narrowed down a microhomology region between the sense *FLAIL* 3′-end in *Arabidopsis* and a non-coding DNA region near the *Camelina* gene *Csa17g011930.* In turn this region in *Camelina* is syntenic to a genomic region in *Arabidopsis* surrounding the *At1G08480* locus (Fig. S3A). Indeed, we successfully detected the existence of genomic *FLAIL-like* non-coding DNA by PCR amplification (Fig. S3C). We next performed RT-PCR targeting the *FLAIL-like* non-coding DNA region to examine RNA expression in *Camelina sativa* leaves and seedlings (Fig. S3B-C). Our data are consistent with the expression of RNA from the *FLAIL-like* non-coding DNA region in *Camelina sativa*, even though we detected noticeably weaker expression compared to *Arabidopsis FLAIL* expression in equivalent experimental conditions. In conclusion, our comparative genomic analysis identified a candidate *FLAIL-like* non-coding DNA region in Brassicaceaes with signatures of RNA expression.

### *FLAIL* characterizes a *trans-*acting lncRNA repressing flowering

To address the function of *FLAIL* in *Arabidopsis*, we used CRISPR/Cas9 technology with paired sgRNAs to generate two different *flail* knockout mutants with deletion fragments of 229 bp (*flail1*) [44], and 343 bp (*flail2*) (Fig. S4A-E). We also obtained a mutant line SAIL_645_C03 (*flail3)* carrying a T-DNA insertion at the 3′-end of sense *FLAIL* locus (Fig. S4A, F). All three mutants reduced the expression of sense and antisense *FLAIL* transcripts as detected by RT-qPCR, suggesting that they reduced the bioavailability of *FLAIL* isoforms (Fig. 1C-D). All three *flail* mutants flowered earlier than wild type (Fig. 1E-F). Our genetic data thus revealed a link between non-coding transcription at the *FLAIL* locus and flowering time.

To test which transcript of *FLAIL* was linked to the observed phenotype of the *flail* mutant we performed a complementation test. We transformed a DNA fragment encoding either sense or antisense *FLAIL* driven by the corresponding native promoter into the *flail3* mutant background with *GUS* driven by *35S* promoter as a control (Fig. 2A). Importantly, the constructs carried the *NOS* terminator that largely abolished initiation of antisense transcription [8, 37, 45]. We selected *flail3* for complementation because *FLAIL* disruption by T-DNA insertion at the 3′-end left potential sORF regions at the 5′-end intact (Fig. S4A). We selected two representative homozygous single locus insertion lines for each transformed construct (*flail3 pFLAIL:gFLAIL18/88* and *flail3 pasFLAIL:gasFLAIL18/39*) that expressed sense and antisense *FLAIL* at levels slightly higher than or similar to wild type (Fig. 2B-C). However, only exogenous expression of sense *FLAIL* rescued the early flowering phenotype of the *flail3* mutant (Fig. 2D-E), suggesting that the early flowering of the *flail* mutant was caused by the disruption of the sense *FLAIL* transcript isoform. In *Arabidopsis*, transgene insertion is non-targeted. We thus reasoned that complementation argued for the capability of sense *FLAIL* to act from a different genomic location, presumably as a *trans-*acting lncRNA. This hypothesis predicts that knock-down of sense *FLAIL* RNA should result in equivalent effects as genomic mutations. To test this hypothesis, we employed strand-specific RNA repression using artificial microRNAs (amiRNAs) [46]. We generated two amiRNA targets (*amiR-FLAIL-11* and *amiR-FLAIL-12*) (Additional file1: Fig. S5A), both exhibiting strongly reduced sense *FLAIL* transcription (Additional file1: Fig. S5B) and similar effects on flowering as genomic mutations (Additional file1: Fig. S5C-D). In conclusion, our experimental data indicate that the sense *FLAIL* lncRNA represses flowering through a *trans*-acting mechanism.

**Fig. 2.**
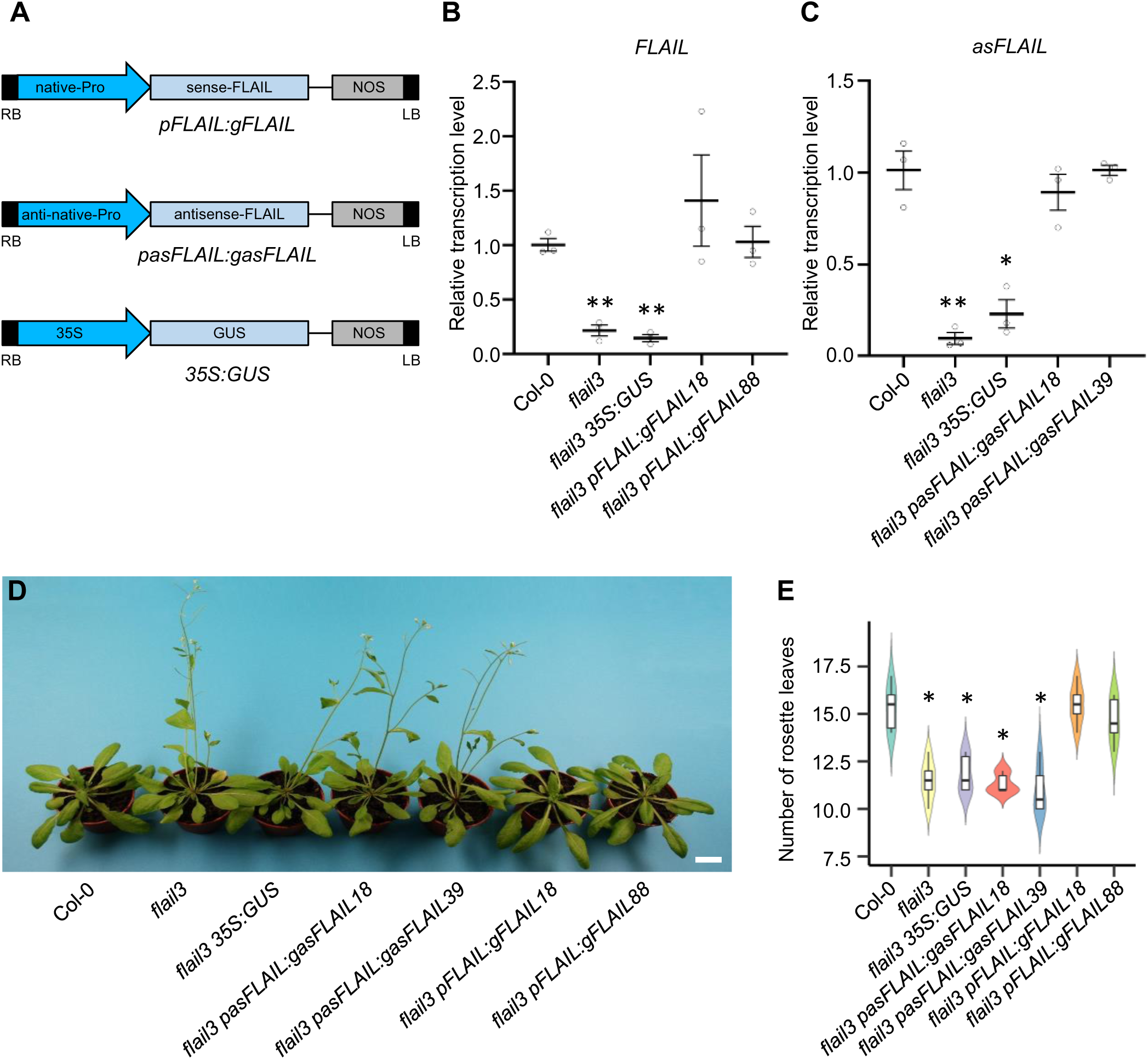
*Sense-FLAIL* RNA is functional for flowering. **A** Schematic representations of the T-DNA constructs containing the native promoter of *sense-FLAIL* (native-Pro) fused to the *sense-FLAIL* DNA region with NOS as a terminator or the native promoter of *anti-sense FLAIL* (anti-native-Pro) fused to the *anti-sense FLAIL* DNA region or a negative control with the 35S promoter (35S) fused to GUS reporter were transformed into *flail3 Arabidopsis* plants. **B-C** Detection of *FLAIL* and *asFLAIL* genes expression in Col-0, *flail3* mutant and complemented lines expressing *pFLAIL:gFLAIL* and *pasFLAIL:gasFLAIL* by RT-qPCR. Transcript levels were normalized to *UBQ* expression levels. Y-axis showed relative values compared to the expression level of Col-0. Error bars represented s.e.m (n = 3 independent 14-d seedling pools). *, *p* value < 0.05 and **, *p* value < 0.01 by Student’s t-test compared to Col-0. **D** Representative morphological phenotypes of 4-week-old plants of Col-0, *flail3* mutant, transgenic lines at 20 °C under a 16-h light/8-h dark growth condition. Scale bar: 2 cm. **E** Violin graph showed number of rosette leaves after appearance of the first flower bud in Col-0. Data represented the mean of six independent experiments. Boxes spanned the first to third quartile, bold black lines indicated median value for each group and whiskers represented the minimum and maximum values. *, *p* value < 0.05 was indicated by Student’s t-test compared to Col-0.

### *FLAIL* lncRNA binding to chromatin regions promotes the expression of selected flowering repressors

Early flowering in *Arabidopsis* may be associated with altered expression of flowering-related genes. To gain insight into the molecular basis of *FLAIL*-mediated regulation of flowering time, we determined the transcriptional profiles of two-week old seedlings for wild type, *flail3,* and *flail3 pFLAIL:gFLAIL* with at least two independent replicates using stranded RNA-seq. Since the early flowering time effect in *flail3* could be rescued by *pFLAIL:gFLAIL*, we reasoned that this experimental setup may be suitable to identify gene expression changes directly correlated with sense *FLAIL* expression. Compared to wild type, 1221 differentially expressed genes (DEGs) were called by DESeq2 with at least two-fold change (adjusted *p* value < 0.05; Table S1), with 419 up-regulated and 802 down-regulated genes in *flail3* mutants. Almost half of these transcriptional differences in *flail3* reverted to wild-type expression level by exogenous expression of the *FLAIL* sense RNA into *flail3* mutants, including the *FLAIL* lncRNA itself (Fig. S6A and Fig. S7F, Table S2). We next focused on functional annotations of genes associated with the process of flowering [47]. Among the DEGs in *flail3*, we identified twenty genes linked to flowering (Fig. S6A). Expression of most of them were fully (eight) or partially (five) rescued by the expression of *pFLAIL:gFLAIL* (Fig. 3A-E, Fig. 4E-H, Fig. S6B, S7A-E and S9B, D). In conclusion, our transcriptomic data indicate potential targets for the *trans-*acting lncRNA *FLAIL*.

**Fig. 3.**
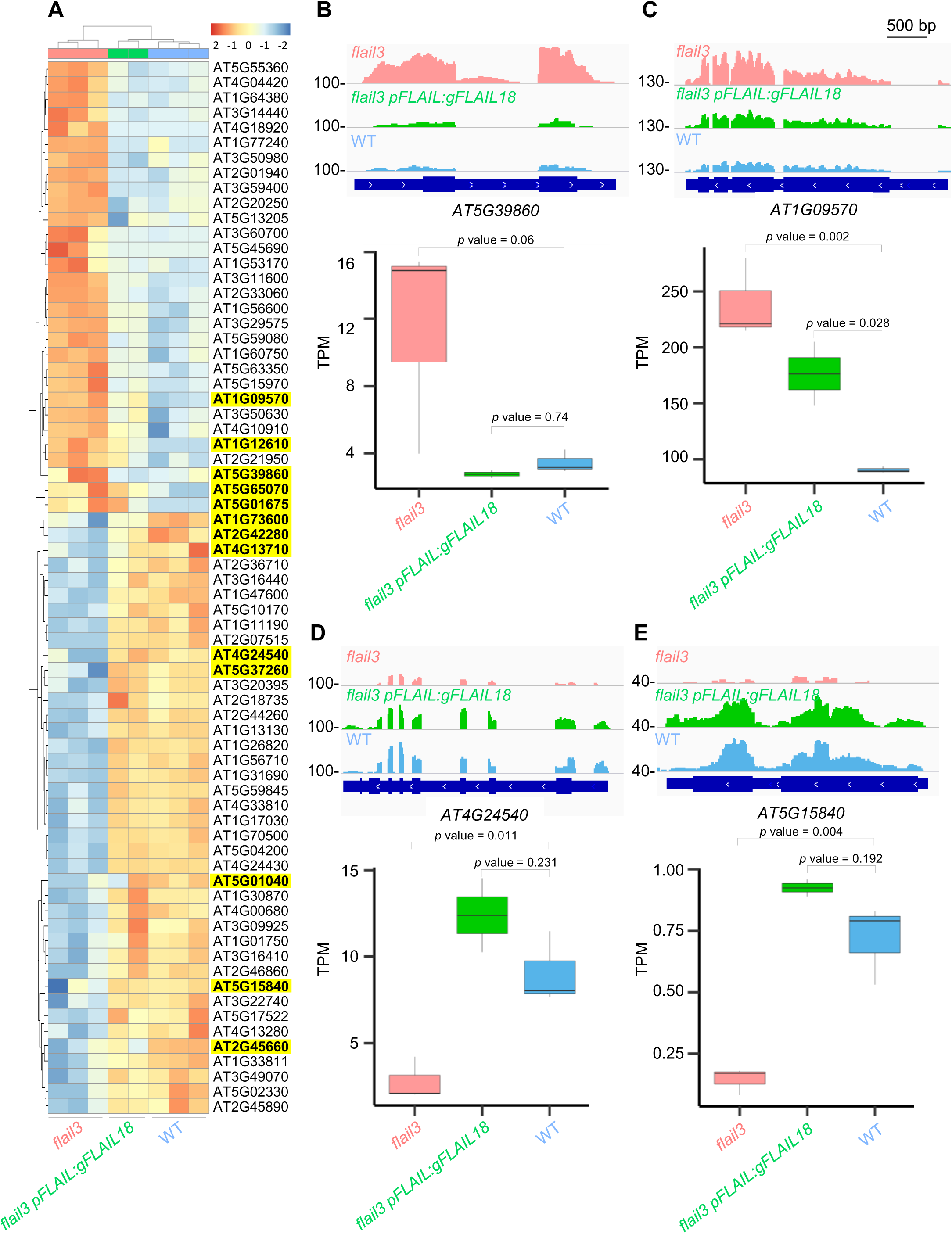
*FLAIL* regulates flowering related genes. **A** Heatmap and hierarchical clustering of top 70 differentially expressed genes (DEGs) in *flail3 versus* Col-0 that were best rescued in the *flail3 pFLAIL:gFLAIL18* complementation line. DEGs analyzed by DESeq2 with |log2 fold change| > 1 and adjusted *p* value < 0.05 were considered significant differential expression. Three biological replicates for *flail3*, wild type, and two for *flail3 pFLAIL:gFLAIL18.* Flowering genes were highlighted in yellow bold. Samples clustered together on the basis of corresponding similar expression profiles. The color scale reflected the log 2-fold change in gene expression, ranging from down-regulated (blue) to up-regulated (red). **B-E** Genome browser screenshots illustrating the expression of dysregulated flowering genes in *flail3* were rescued in complementation line (top panel). Normalized read counts (TPM from RNA-seq) for differentially expressed flowering genes in WT, *flail3*, and *flail3 pFLAIL:gFLAIL18* plants (bottom panel). Boxes spanned the first to third quartile, bold black lines indicated median value for each group and whiskers represented the minimum and maximum values. All *p* values were denoted by Students’ t-test. Bar = 500 bp.

**Fig. 4.**
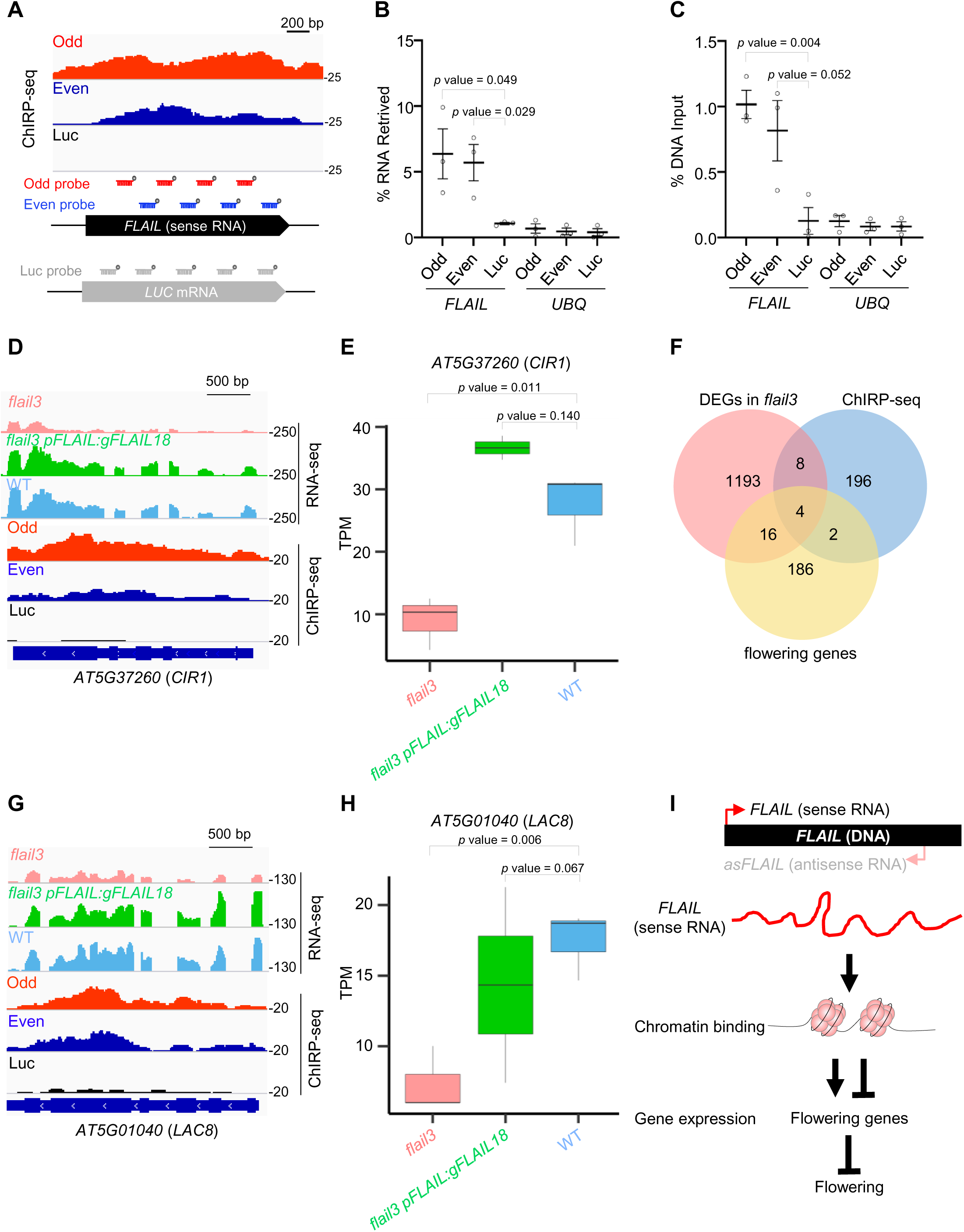
*FLAIL* affects flowering by chromatin binding of flowering genes. **A** Top, *FLAIL* bound the locus itself by ChIRP-seq from two independent Odd and Even probed chromatins. Luc probe was used as a control. Bottom, schematic representation of the antisense oligonucleotide probes that were biotinylated at the 3′-end with Odd (in red) and Even (in dark blue) against *FLAIL* sense RNA and Luc probe (in grey) against *LUC* mRNA. **B** ChIRP-qPCR using probe pools *FLAIL*-asDNA (Odd and Even) retrieved ∼5%–10% of *FLAIL* endogenous RNA and < 1% levels of *UBQ*. *Luc-*asDNA probes retrieved much lower levels of both RNAs as a control. **C** *FLAIL* DNA signal was identified in both Odd and Even probes. *UBQ* region showed much less binding signal in all probes as a negative control. Graphs in **B** & **C** showed the mean ± s.e.m. (n= 3 independent replicates). **D, G** Genome browser screenshots illustrating two of *FLAIL* bound targets *CIR1* (**D**) and *LAC8* (**G**) in RNA-seq (lane 1-3) and ChIRP-seq (lane 4-6). FLAIL binding peaks were called by CCAT3.0 with the cutoff FDR = 0.232. **E, H** Normalized read counts (TPM from RNA-seq) for differentially expressed (DE) flowering genes in WT, *flail3*, and *flail3 pFLAIL:gFLAIL18* plants (bottom panel). Boxes spanned the first to third quartile, bold black lines indicated median value for each group and whiskers represented the minimum and maximum values. **F** Venn diagram of genes targeted by *FLAIL* (ChIRP-seq) and genes overlapped with DEGs in *flail3* and flowering related genes. **I** A model for the *trans-*acting *FLAIL* sense RNA regulated flowering. The *FLAIL* sense RNA binds chromatin regions of flowering genes to regulate expression levels of flowering related genes and thus affects flowering time. All *p* values were denoted by Students’ t-test.

The reversible effect on flowering gene expression upon re-introduction of *pFLAIL:gFLAIL* argued for a direct effect of the *FLAIL* lncRNA. To identify where *FLAIL* bound chromatin regions, we performed chromatin isolation by RNA purification (ChIRP) of endogenous *FLAIL* followed by DNA-seq (*FLAIL* ChIRP-seq). We targeted sense *FLAIL* by two non-overlapping antisense oligonucleotide pools (Even and Odd probes, Fig. 4A bottom) tiled along the entire *FLAIL* transcript sequence compared to an oligonucleotide pool against the *LUCIFERASE* (*LUC*) mRNA as control that is not expressed in our strains. We efficiently captured the endogenous *FLAIL* RNA from chromatin compared to the unrelated *UBQ* mRNA (Fig. 4B). Moreover, *LUC*-specific probes did not enrich *FLAIL*, suggesting the specificity of RNA affinity purification in our assay conditions (Fig. 4B). After isolating DNA fragments associated with the *FLAIL*-containing complex, we assessed the genome-wide occupancy of *FLAIL* at high resolution by ChIRP-seq, followed by peak calling using CCAT3.0 [48]. Only peaks that occurred at target genes from both even and odd probe pools, but not from two or more independent experiments of Luc pools were considered significantly enriched (Fig. S8A). This analysis strategy allowed us to identify 210 target genes of *FLAIL* (Table S3). We observed a strong enrichment for the *FLAIL* locus by both ChIRP-seq (Fig. 4A, top) and ChIRP-qPCR (Fig. 4C). We noted that the *FLAIL* locus represented the only locus where we could expect binding, thus validating our identification strategy of genomic *FLAIL* binding. We analyzed the overlap between genes which were differentially expressed in the RNA-seq experiment and those genes that showed statistically significant *FLAIL* binding by ChIRP-seq (Fig. 4F). We identified twelve genes matching both criteria. At these targets, chromatin binding by *FLAIL* was linked to corresponding changes of gene expression, arguing for direct effects of *FLAIL* binding on gene expression. Four of these genes were functionally connected to flowering: *CIRCADIAN 1 (CIR1), LACCASE 8 (LAC8), PECTIN LYASE-LIKE 25 (PLL25) and PHOSPHOETHANOLAMINE N-METHYLTRANSFERASE 3 (NMT3)* (Table S4). All four flowering genes were transcriptionally down-regulated in *flail3* (Fig. 4D-E and 4G-H, Fig. S9A-D, Table S1).

However, the complementation construct failed to restore *NMT3* expression, arguing against a direct effect here (Fig. S9A-B, Table S2). A detailed examination revealed that *PLL25* regulates flower morphology rather than flowering time [49], arguing against effects on flowering time through mis-regulation of this gene [50, 51]. On the other hand, *cir1* [52] and *lac8* [53] mutants displayed early flowering phenotypes. These data suggest repression of *CIR1* and *LAC8* as candidate molecular hypothesis to explain the early flowering time phenotype in *flail* [52, 53]. In conclusion, our data suggest that *FLAIL* sense RNA represses flowering through inhibiting expression of the direct *FLAIL* targets *CIR1* and *LAC8*, consistent with a model where gene expression changes regulating flowering are mediated by interactions of the *trans-*acting lncRNA *FLAIL* with the genome (Fig. 4I).

## Discussion

### *FLAIL* functions in flowering time

In this study, we report repression of flowering time by the *trans-*acting lncRNA *FLAIL*. While many lncRNAs affect flowering regulation, the molecular mechanisms remain largely unclear [15–20]. Previously characterized lncRNAs including *COOLAIR*, *COLDAIR*, *ASL*, *COLDWRAP*, and *MAS* are transcribed from the flowering genes *FLC* or *MAF4* to locally affect their coding gene expression in *cis* [20, 24, 27, 54, 55]. We note that the *FLAIL* 3′-end resides approximately 130 bp upstream of the 5’ UTR region of the *PORCUPINE* (*PCP*, also called *Sm protein E1*, *SME1*) gene, and *pcp/sme1* mutants flower early [56, 57]. We failed to find evidence for the hypothesis that *FLAIL* may affect flowering through *PCP* by a *cis*-acting mechanism. *FLAIL* neither regulates *PCP* expression nor shows overlapping effects on gene expression (Table S1 and S5-7, Fig. S10A-D).

In contrast, our work identifies a *trans-*acting *FLAIL* that represses flowering and binds multiple genomic loci. We thus favor the interpretation that *FLAIL* binding to other genomic regions explains the phenotypic effects of the mutants. For instance, *FLAIL* promotes the expression of the MYB family transcription factor *CIR1*, which in turn broadly contributes to gene expression. This may also explain why some genes are *FLAIL*-regulated but lack chromatin interactions with *FLAIL*. *CIR1* is a circadian clock gene, induced by light and involved in a regulatory feedback loop that controls a subset of the circadian outputs [52]. Our GO analysis supports that a subset of DEGs are connected to the response to red or far red light that contains among other key flowering genes such as *PHYTOCHROME INTERACTING FACTOR 4* (*PIF4*) and *CONSTANS* (*CO*) (Fig. S10B). *FLAIL* also binds the chromatin region of *LAC8*. LAC8 is a laccase family member that mainly modulates phenylpropanoid pathway for lignin biosynthesis [53, 58]. Similar to *flail*, *lac8* mutants flower early [53]. While intermediates in this pathway [59, 60] or dysregulation of lignin-related genes [21, 22, 61] could promote flowering in plants, the molecular connections of reduced *LAC8* expression to effects on flowering time will require further investigation. Nevertheless, our observations of reduced expression of *LAC8* and *CIR1* in *flail*, combined with restored expression upon re-introduction of *FLAIL*, and direct *FLAIL* binding to *LAC8* and *CIR1* chromatin, suggested that early flowering in *flail* may result from combined direct and indirect effects of *LAC8* and *CIR1* repression.

The range of plausible candidate mechanisms by which *FLAIL* may promote gene expression includes targeting of chromatin modifying complexes to activate gene expression as shown for the mammalian lncRNA *HOTTIP* [62], or effects on pre-mRNA RNA processing to stimulate mRNA expression as indicated for the plant lncRNA *ASCO* [63–65]. While our results clarify key aspects of functional units of the non-coding genome, it will remain an exciting future research endeavor to elucidate how sense *FLAIL* mediates the activation of floral repressor genes at the molecular level.

### Identification of the functional *FLAIL* RNA isoform of the *FLAIL* locus

A challenge in the field of lncRNA biology is the functional characterization of lncRNA loci. Our functional dissection of *FLAIL* illustrates these challenges, yet reveals a compelling example for a *trans-*acting lncRNA. Like many other loci, *FLAIL* generates multiple transcript isoforms, including cryptic transcript isoforms that would be missed by most standard transcriptomic approaches. DNA regulatory elements embedded in lncRNA loci and the act of lncRNA transcription may regulate the expression of neighboring genes (*cis-*acting) [66, 67]. However, disruption of the *FLAIL* locus had no significant impact on gene expression in a surrounding ∼150 kb genomic window (from upstream AT2G18560 to downstream AT2G18940). These observations argue against the hypothesis that the *FLAIL* locus acts as a *cis-*acting RNA or DNA element. In contrast, we found genetic evidence that the *FLAIL* sense lncRNA functions as a *trans-*acting RNA. First, exogenous expression of *FLAIL* sense RNA *in vivo* rescues the early flowering phenotype as well as the expression of flowering genes in *flail3.* The rescue is specific for the *FLAIL* lncRNA encoded by the sense strand. Second, amiRNA mutants that specifically knock-down *FLAIL* sense RNA levels without effect on the DNA also show early flowering. Collectively, these data indicate that the role of the *FLAIL* locus in the repression of flowering is executed by a *trans-*acting lncRNA derived from the sense strand.

We assessed the protein-coding potential of *FLAIL* using a series of software tools that give low scores, arguing against a protein encoded by *FLAIL*. Nevertheless, Ribo-seq identifies ribosome association of *FLAIL,* consistent with two small ORFs with ∼9 amino acids (Fig. S1B) [39]. Even though we found no evidence for peptide production in proteomics data, sORFs embedded in lncRNAs may encode for functional peptides [68, 69]. In our allelic series of *flail* mutants only *flail1* mutates the potential sORFs, while *flail2* and *flail3* keep the DNA sequences encoding the potential sORFs intact. The range of genetic mutations thus argue against the contribution of potential sORFs peptides to flowering, since the early flowering phenotype is similar in all three mutant backgrounds. Moreover, the microhomology with *Camelina* mapped to the 3’-region of *FLAIL* where we identified no ribosome association. While future experimental research in *Camelina sativa* would be needed to examine functional conservation in flowering, the microhomology of *FLAIL* in other Brassicaceae is consistent with functional RNA domains that are a characteristic of conserved *trans*-acting lncRNAs.

## Conclusions

In summary, this work highlights the contribution of lncRNAs to fine tune the complex developmental transition to initiate flowering. Regulation for key developmental decisions by lncRNA-based mechanisms may confer specific organismal advantages, yet detecting the functional roles of the non-coding genome may also be facilitated in the developmental context. RNA-based regulation of plant reproductive development through *trans-*acting lncRNA promises future possibilities for plant breeding research to improve Brassicaceae crop quality and resilience.

## Methods

### Plant materials and growth conditions

Table S8 provides a complete overview of the plant materials used in this study. Seeds of *flail3* (SMA2648, Table S8) were obtained from the Nottingham *Arabidopsis* Stock Centre and genotyping was performed using primers Mlo37/1788/1789 (Additional file 2 Table S9). *A. thaliana* plants were grown in a growth chamber with a long day photoperiod (16-h light/8-h dark) at 20 °C at a photo flux density of approximately 100 μmol/m^2^/sec in the chamber. For seedling treatments, seeds were surface-sterilized and placed on ½ MS + 1% sucrose plates at 4 °C in darkness for 3 days prior to germination. Then, plates were placed in the growth chamber in control conditions at 20 °C and sampled.

### CRISPR/Cas9-directed *flail* mutants

Construction of a dual sgRNA-directed *flail1* (*goi*, [44]) knockout SMA3677 (Table S8) was performed by CRISPR/Cas9 using the described protocol [44]. Similarly, to generate the CRISPR mutant *flail2* (SMA3678, Table S8), plasmid pHEE2E-TRI (SMC528) harboring two sets of gBlocks including gBlock1 (a U6-26 promoter, a 19 bp target sequence 1, a sgRNA scaffold, a terminator) and gBlock2 (a U6-29 promoter, a 19 bp target sequence 2, a sgRNA scaffold, a terminator) served as a template to amplify the middle border (BsaI-overhang1-protospacer1-scaffold-terminator-U6-29 promoter-protospacer2-overhang2-BsaI) using primer paris Mlo1757 & Mlo1758 (Table S9). Second, both pKIR1.1 plasmid (SMC529) and the middle border were digested by AarI and BsaI, respectively, then the middle border was integrated into the linearized pKIR1.1 backbone to generate pKIR1.1-dual- sgRNA2 SMC575 (Table S10) for plant transformation. Third, after transformation using *Agrobacterium tumefaciens* strain SMA111 (Table S11), the T1 generation plants for successful T-DNA insertion events were selected by identifying seeds with red fluorescence, then seeds from individual T1-genotyped plants were harvested and grown. Finally, T2 mutants that did not contain the CRISPR/Cas9 construct were identified by picking up T2 seeds without red fluorescence. PCR products were amplified from Cas9-free T2 plants using oligonucleotides Mlo2478/2479 (Table S9) flanking the deletion site for Sanger sequencing.

### Complementation assay

Complementation constructs were generated using SMC431. *pFLAIL:FLAIL* and *pasFLAIL:asFLAIL* were amplified from genomic wild type DNA using primers MLO1746/1747 and MLO1759/1761, respectively (Table S9). The resulting PCR products were inserted into pENTR-D-Topo by topo cloning to generate entry vectors SMC542 and SMC541 (Table S10). The entry vectors were used in a LR reaction with SMC431 to generate expression vector SMC546 (containing *pFLAIL:gFLAIL* construct) and SMC545 (containing *pasFLAIL:gasFLAIL* construct) (Table S10). The complementation constructs together with the control vector *35S:GUS* (SMC377) were then transformed into GV3101 to get strains SMA112-114 (Table S11), followed by transformation into the *flail3* mutant. Seeds from transformed *Arabidopsis* plants were screened for T-DNA integration by hygromycin resistance. Multiple independent single-locus insertions were identified by segregation analysis and homozygous T3 transgenic plants SMA4477 for *flail3 35S:GUS*, SMA4462 for *flail3 pFLAIL:gFLAIL18*, SMA4464 for *flail3 pFLAIL:gFLAIL88*, SMA4467 for *flail3 pasFLAIL:gasFLAIL18*, SMA4468 for *flail3 pasFLAIL:gasFLAIL39* were used for the complementation assay (Table S8).

### Cloning of amiRNA

To construct *amiR-FLAILs*, we designed two artificial miRNA sequences in the miR319a backbone using Web MicroRNA Designer (WMD3) software to generate four oligonucleotide sequences (I to IV) (Mlo1774-1781, Table S9), which were used to engineer the amiRNA into the endogenous miR319a precursor by site-directed mutagenesis. The amiRNA containing precursor was generated by overlapping PCR using SMC532 as a template following the protocol described in the WMD3 publication [46]. The fragment containing the amiRNA sequence was then introduced into pENTR-D-Topo by topo cloning to generate entry vectors (SMC547 for *amiR-FLAIL11-Topo* and SMC548 for *amiR-FLAIL12-Topo*) (Table S10). The entry vectors were used in a LR reaction with SMC531 to generate expression vector SMC558 (containing *amiR-FLAIL11* construct) and SMC559 (containing *amiR-FLAIL12* construct). Plant transformation was performed using *Agrobacterium* strains SMA115-117 (Table S11). Finally, homozygous T3 transgenic plants SMA4469, SMA4471, and SMA4480 (Table S8) were used in this study.

### Gene expression analysis by PCR/RT-(q)PCR

For the comparative expression analysis with *C. sativa*, genomic DNA was extracted from wild type *A. thaliana* and *C. sativa* seedlings (∼1.5 weeks) and mature leaves (∼4 weeks) using the DNeasy Plant Mini kit (Qiagen #69104) and diluted to 5 ng/μl. Total RNA was extracted from the same tissues using the RNeasy Plant Mini kit (Qiagen #74904) with RNase-free DNase set (Qiagen #79254). Reverse transcription (RT) was performed on 1 μg of DNase-treated total RNA using the iScript cDNA synthesis kit (Bio-Rad #1708890), with the same amount of total RNA diluted in water as negative controls for RT-PCR. Primers (Table S9) were designed to amplify *AthaFLAIL* (*AT2G18735*), *CsatFLAIL-like* (*C. sativa*, Ensembl v51 Chr17 3432317-3432814), and *GAPDH* as an amplification control. Two reverse primers were designed to target different portions of *AthaFLAIL* and *CsatFLAIL-like*. The *GAPDH* primers amplify both *AthaGAPDH* and *CsatGAPDH*. PCR was performed with templates consisting of either 20 ng of genomic DNA, 50 ng of total RNA, or 50 ng of total RNA that had been reverse transcribed. 35 cycles of amplification were performed for all template types. PCR products were visualized after agarose gel electrophoresis using the Bio-Rad Chemi-doc imaging platform.

For reverse transcription quantitative real-time PCR, total RNA was extracted from two-week old *Arabidopsis* seedlings using an RNeasy Plant Mini Kit (Qiagen, Germany). DNA in the isolated RNA were digested with TURBO DNase (Thermo Fisher Scientific, USA). Purified RNA was subsequently reverse transcribed into cDNA with iScript™ cDNA Synthesis Kit (Bio-Rad, USA) following manufacturer’s instructions. For real-time PCR analysis, the resulting cDNA was diluted ten-fold and used as a template in a PCR reaction with GoTaq qPCR Master mix (Promega, USA) and run on a CFX384 Touch instrument (Bio-Rad, USA) with an initial denaturation at 95 °C for 2 minutes, followed by 40 cycles at 95 °C for 15 seconds, 60 °C for 1 minute. Primer efficiencies were evaluated on a standard curve generated using a 10-fold dilution series of the sample over four dilution points. Relative expression was calculated and normalized to the internal reference gene *UBQ*. All primers (Table S9) did not show any evidence for non-specific products in the melting curve analysis.

### Chromatin isolation by RNA purification (ChIRP)

ChIRP was performed as previously described with some modifications [70]. Probe design: The antisense oligonucleotide probes were designed against the full-length *FLAIL* sequence using the online probe designer at www.singlemoleculefish.com [71]. The probes were biotinylated at the 3′ end. To assess the specificity of the target capture by the oligonucleotides, the *FLAIL* probes were divided into two pools, Odd (Mlo2885) and Even (Mlo2886). All the experiments were carried out using both pools independently. Sixteen biotinylated oligonucleotide probes complementary to the *LUCIFERASE* transcript were pooled (Mlo2887) as negative control (Table S9).

RNA immunoprecipitation: Two-week old seedlings were crosslinked in 3% formaldehyde solution by vacuum infiltration for 25 minutes and then quenched by the addition of 0.125 M glycine at room temperature for 5 minutes. After washing and drying, crosslinked seedlings (2.5 g) were ground to a fine power in liquid nitrogen and suspended in 10 ml Honda Buffer (0.44 M Sucrose, 1.25% Ficoll, 2.5% Dextran T40, 20 mM Hepes KOH with pH 7.4, 10 mM MgCl_2_, 0.5% Triton X-100, 5 mM DTT, 1 mM PMSF, 1 × Cocktail, and 2 U/ml RNase inhibitor). The samples were put on ice and mixed gently at 4 °C until the solution became homogenous, then filtered through two layers of Miracloth, centrifuged at 4000 rpm for 15 minutes at 4 °C. Pellets were resuspended in 1.5 ml Honda buffer and centrifuged at 4000 rpm for 15 minutes at 4 °C. Resuspending and centrifugation were repeated until the pellet was no longer green, typically two more times (∼15 minutes). Pellets were suspended in 1600 μl nuclear lysis buffer (50 mM Tris-HCl, pH 7.5, 2 mM MgCl_2_, 1% SDS, 0.1 mM PMSF, 10 mM EDTA, 1 mM DTT, 1 × Cocktail, 0.1 U/μl RNase Inhibitor) and sonicated at M setting 5 minutes (30 seconds on and 30 seconds off) × 3 times by sonicator (QSONICA Q700, USA) until DNA was fragmented into 200–500 bp pieces. After centrifugation at 12,000 g at 4 °C for 10 minutes, the supernatant (around 1.5 ml) was diluted with 2 volumes of pre-heated hybridization buffer (750 mM NaCl, 1% SDS, 50 mM Tris, pH 7.5, 1 mM EDTA, 15% formamide, and 1 × protease inhibitor). The clear mixture was divided into 4 equal aliquots (IP-RNA and IP-DNA for both Even and Odd probes). Biotinylated DNA probe (2 µl of 100 pmol/μl) was added to each aliquot and incubated at 37 °C for 4 hours with gentle mixing. Then 50 µl of well-washed Streptavidin C1 magnetic beads were added to each sample and incubated at 37 °C for 30 minutes. Captured beads were washed three times with high salt wash buffer (2 × SSC, 0.5% SDS, 1 mM DTT, and 1 mM PMSF) and three times with low salt wash buffer (0.1 × SSC, 0.5% SDS, 1 mM DTT. and 1 mM PMSF).

RNA isolation: Beads were resuspended in 200 µl RNA elution buffer (100 mM NaCl, 50 mM Tris-HCl with pH 7.5, 1 mM EDTA, 1% SDS) and boiled for 15 minutes. RNA samples were treated with Proteinase K (1 mg/ml) at 65 °C for 1 h while shaking. RNA was extracted using TRIzol method, and then treated with DNase (Qiagen). RNA was used for RT-qPCR analysis to confirm RNA retrieval.

DNA isolation: DNA was eluted with elution buffer (200 mM NaCl, 50 mM NaHCO_3_, 1% SDS, 10% SDS, 0.1 U/µl RNase H). After Proteinase K treatment at 45 °C for 1 h, DNA was extracted using ChIP DNA Clean & Concentrator (Zymo Research) and used for subsequent qPCR analysis or high-throughput sequencing.

### Comparative Genomics

FLAIL sequence similarities were initially identified through reciprocal best BLAST against eleven disparate Brassicaceae genomes in CoGe BLAST [72] with an e-value of 10^-5^. As no hits were identified in a species more distantly related than *Camelina sativa*, comparative analysis was restricted to *Arabidopsis* and *Camelina*. Genomic regions displaying sequence similarity to the *Arabidopsis FLAIL* locus in the *Arabidopsis thaliana* (TAIR10) and *Camelina sativa* (Ensembl v2.0) were identified using CoGe BLAST, with regions of microsynteny identified using GEvo [73]. Nucleotide sequence for syntenic regions were extracted and imported into Geneious Prime v2021.2.2 [74]. Multiple sequence alignments were used to design primers that specifically amplified a portion of *FLAIL* and the *FLAIL-like* intergenic region in *Camelina*.

### Statistical analysis

The number of rosette leaves were measured with ImageJ software [75]). Statistical analysis was performed using the R software [76] or GraphPad Prism 9 [77]. For comparisons, data were evaluated using Student’s t-test for significance of differences: * means *p* value < 0.05, ** means *p* value < 0.01.

### Bioinformatics

All the supporting code for bioinformatics analysis is repository available at FLAIL_2021 from GitHub (https://github.com/Yu-Jin-KU) [78].

### Sequencing analysis

RNA-seq libraries were prepared using NEXTFLEX® Rapid Directional RNA-seq Library Prep Kit (NOVA-5138-08) and ChIRP-seq libraries were constructed using the ChIP-seq Library Prep Kit (NOVA-5143-02 with NEXTflex® ChIP-seq Barcodes-24) following the manufacturer’s protocol and quantified on Agilent 2100 Bioanalyzer. Both RNA-seq and ChIRP-seq libraries were pooled into one flow cell (NextSeq 500/550 High Output Kit v2.5) for sequencing in PE mode (2*75 bp) on Illumina Nextseq 500.

Previously published RNA-seq datasets for wild type and PCP (SME1) deficient plants were downloaded from European Nucleotide Archive (accession number PRJEB24412) and from Gene Expression Omnibus (accession number GSE116964). The raw reads were quality controlled by the FastQC software [79], adapter trimmed by Trimmomatic v0.39 [80] in paired-end mode and then aligned to TAIR10 genome assembly by STAR v2.7.8a [81] in Local mode. Aligned reads with MAPQ below 10 were removed by Samtools v1.1.2 [82]. BAM files were converted to unstranded Bedgraph files using BEDtools genomecov v2.30.0 [78]. The code was detailed in the RNA-seq.sh script on the GitHub page mentioned above. Differentially expressed genes (DEGs) were called using the DESeq2 tool [83]. Expression fold change values were log2 transformed to identify genes with statistically significant differential expression using the following criteria: |log2 fold change| > 1 and adjusted *p* value < 0.05. GO enrichment analysis was done by Metascape [47].

ChIRP-seq reads were quality controlled by the FastQC software [79], adapter trimmed using Trim Galore v0.6.7 [84] and mapped to the TAIR10 genome using STAR v2.7.8a [81]. The converted BAM files were sorted and filtered for MAPQ≥5. After removing PCR duplicates by Samtools v1.1.2 [82], peaks of *FLAIL* occupancy were called with CCAT3.0 [48]. By using CHIPseeker [85] and TxDb.Athaliana.BioMart.plantsmart28 [86] packages in R, peak annotation was performed with the definition for a promoter being 300 bp around the TSS. Alignment statistics were provided in Table S3. Visualization and analysis of genome-wide enrichment profiles were done with IGV genome browser [87].

## Abbreviations

amiRNAs: artificial microRNAs
ChIRP: chromatin isolation by RNA purification
CIR1: CIRCADIAN 1
CNIT: Coding-NonCoding Identifying Tool
CO: CONSTANS
CPC2: Coding Potential Calculator
DEGs: differentially expressed genes
FLAIL: flowering associated intergenic lncRNA
FLC: FLOWERING LOCUS C
LAC8: LACCASE 8
lncRNAs: long non-coding RNAs
LUC: Luciferase
NMT3: PHOSPHOETHANOLAMINE N-METHYLTRANSFERASE 3
sORFs: small open reading frames
PCP: PORCUPINE
PIF4: PHYTOCHROME INTERACTING FACTOR 4
plaNET-seq: plant Native Elongating Transcripts sequencing
PLL25: PECTIN LYASE-LIKE 25
RNAPII: RNA polymerase II
SAMs: shoot apical meristems
SME: Sm protein E1
TIF-seq: Transcript Isoform sequencing
TSS-seq: Transcription Start Site sequencing
UBQ: UBIQUITIN
WMD: Web MicroRNA Designer

## Supplementary information

**Table S1.** DEGs in *flail3* mutant

**Table S2.** DEGs in *flail3 pFLAIL gFLAIL18* plants

**Table S3.** *FLAIL* targets called by CCAT3.0

**Table S4.** Overlapping genes between DEGs in *flail3* and *FLAIL* targets.

**Table S5.** Gene Ontology of DEGs in *flail3*

**Table S6.** Gene Ontology of DEGs in *pcp* (RNA-seq data from Capovilla et al with the ENA accession: PRJEB24412)

**Table S7.** Gene Ontology of DEGs in *sme1* (RNA-seq data from Huertas et al with the GEO accession: GSE116964)

**Table S8.** Seeds

**Table S9.** Primer sequences

**Table S10.** Plasmids

**Table S11.** GV3101 strains

## Declarations

### Ethics approval and consent to participate

Not applicable

### Consent for publication

Not applicable

### Availability of data and materials

The RNA-seq and ChIRP-seq data were deposited at GEO with the number GSE186215 [88]. The reagents described in this study are available from the corresponding author.

### Authors’ contributions

Y.J. and S.M. conceived the study. Y.J. performed most experimental work, A.N.D. and A.D.L.N contributed comparative genomics and expression analysis in S3. M.I. and Y.J. did the computational analysis. Y.J. and M.S. wrote the manuscript.

## Acknowledgements

We thank Prof. Anders H. Lund and for kindly providing Luc probes. We thank Prof. Peter Brodersen for sharing amiRNA vectors. We thank Dr. Pan Zhu and Dr. Marta Montes for assistance with ChIRP, Prof. Egle Kudirkiene and Dr. Quentin Thomas for sequencing. We thank Jasmin Dilgen, Evangelia Lakita, Ida Damholt Richardt, Lei Li for technical assistance, Jan Høstrup for excellent plant care, Dr. Deyong Zhu and the members of Marquardt lab for discussions and manuscript feedback.

---

**Fig. S1.**
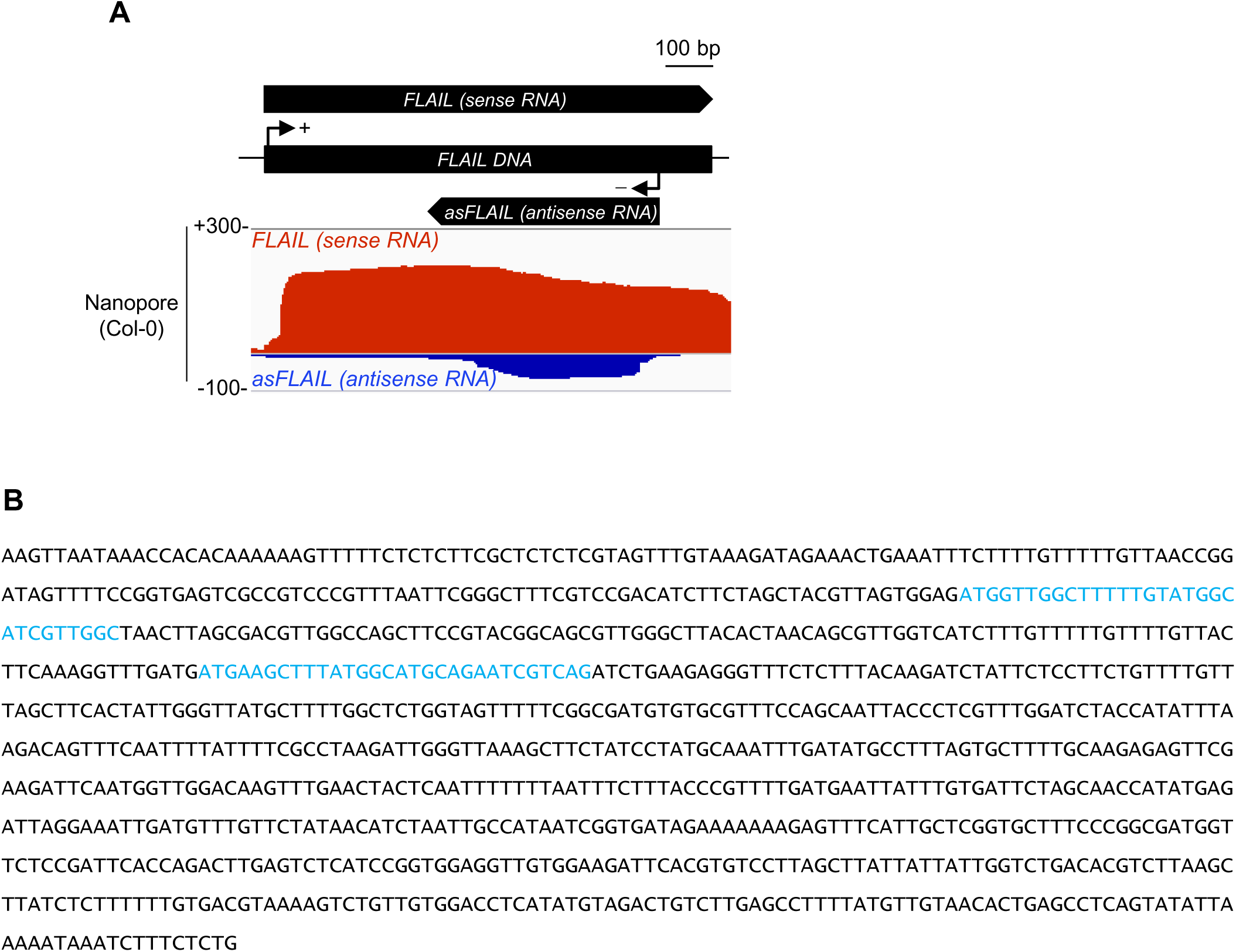
Characterization of *FLAIL* locus. **A** Genome browser view of *FLAIL* splicing status in nanopore sequencing of Col-0. Transcription of sense *FLAIL* RNA and *asFLAIL* RNA were shown in red and dark blue, respectively and no isoform resulting from alternative splicing was observed. **B** *FLAIL* nucleotide sequences in black and two sORFs locations in blue.

**Fig. S2.**
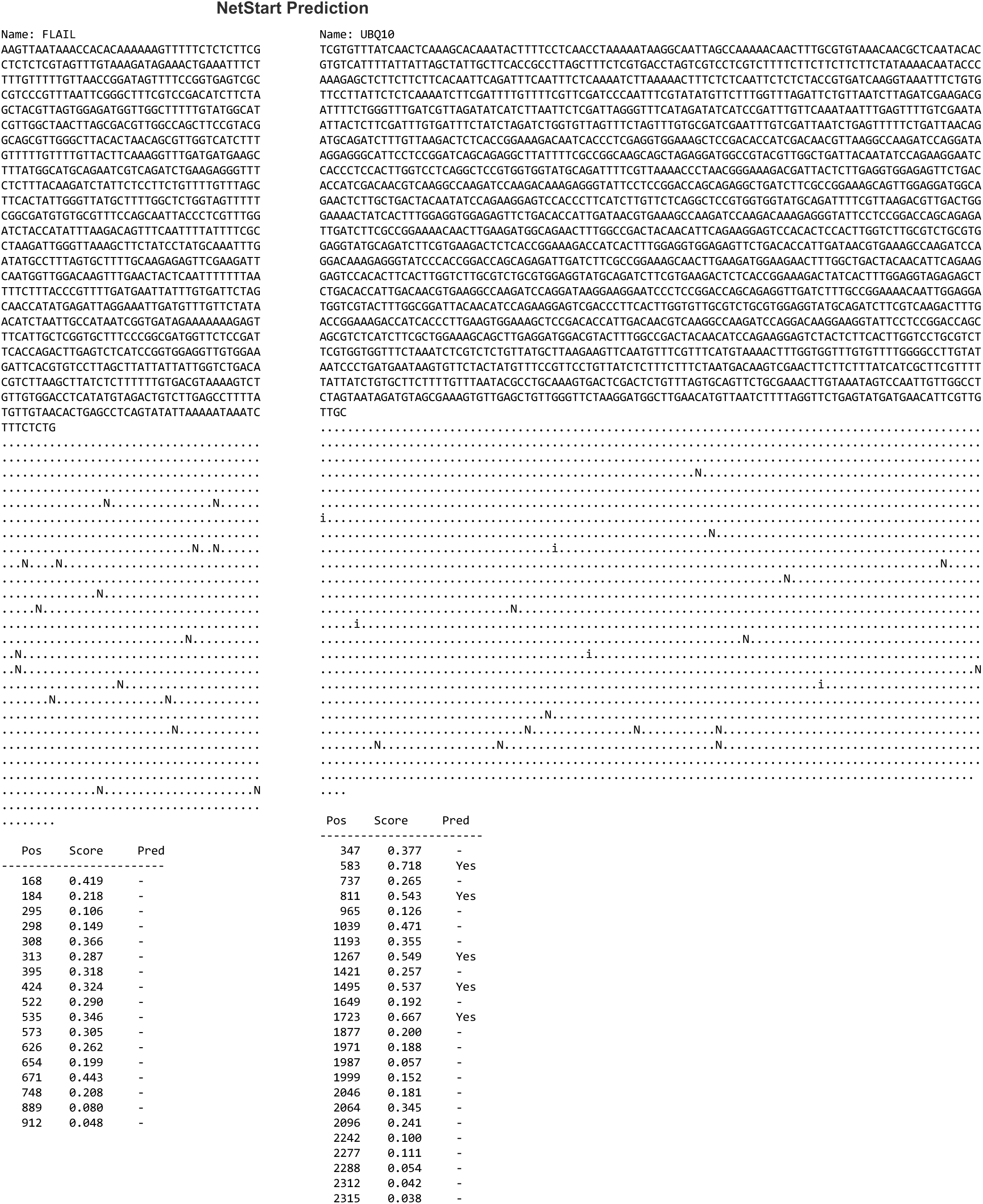
Assessment of *FLAIL* and *UBQ* for protein coding potential. Initiation codon translational analysis using NetStart for *FLAIL* and *UBQ*. The predicted initiation codons were depicted with the letter i, other instances of “ATG” by the letter “N” (non-start). The dots (“.”) were place holders for all the other sequence elements. The scores were always in [0.0, 1.0]; when greater than 0.5, they represented a probable translation start.

**Fig. S3.**
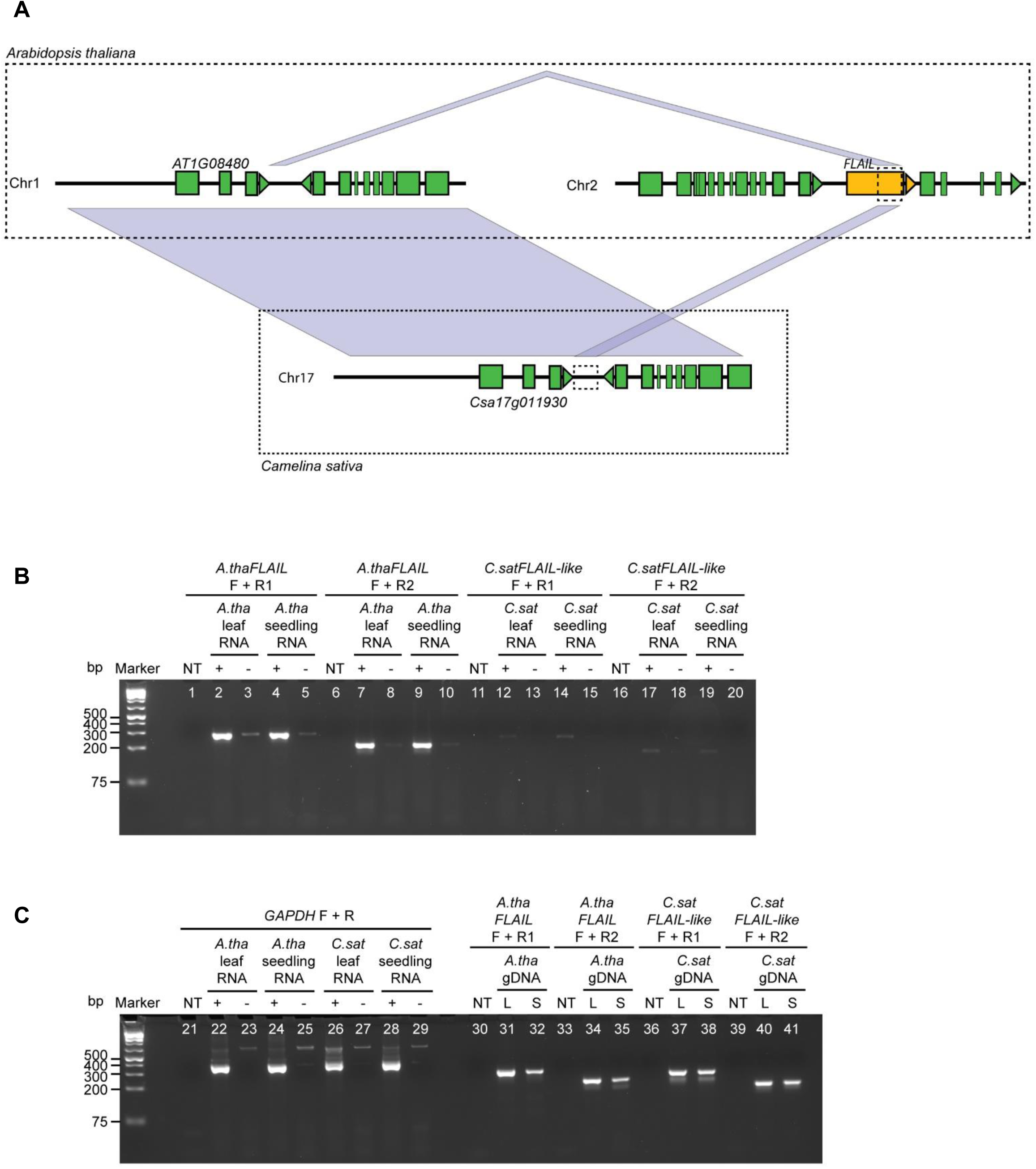
Sequence and transcriptional conservation at the *FLAIL* and *FLAIL-like* loci in *Arabidopsis* and *Camelina*. **A** Schematic depicting the conservation of the *Arabidopsis* (*A.tha*) *FLAIL* locus in *Camelina sativa* (*C.sat*). Green boxes represent exons, with triangles representing direction of transcription. The *FLAIL* locus is represented by a yellow box. Faded blue lines represent sequence similarity between different loci. Dashed boxes at the *FLAIL* and *Csa17g011930* loci represent regions targeted for RT-PCR. **B** Amplification of *A.thaFLAIL* and *C.satFLAIL*-like, using RNA template (+/- RT, lanes 1-20). NT indicates no template was added to the reaction. **C** Control amplifications: *GAPDH* was amplified using RNA template (+/- RT, lanes 21-29), and *A.thaFLAIL* and *C.satFLAIL-like* were amplified using genomic DNA template (lanes 30-41). L indicates leaf tissue, S indicates seedling tissue.

**Fig. S4.**
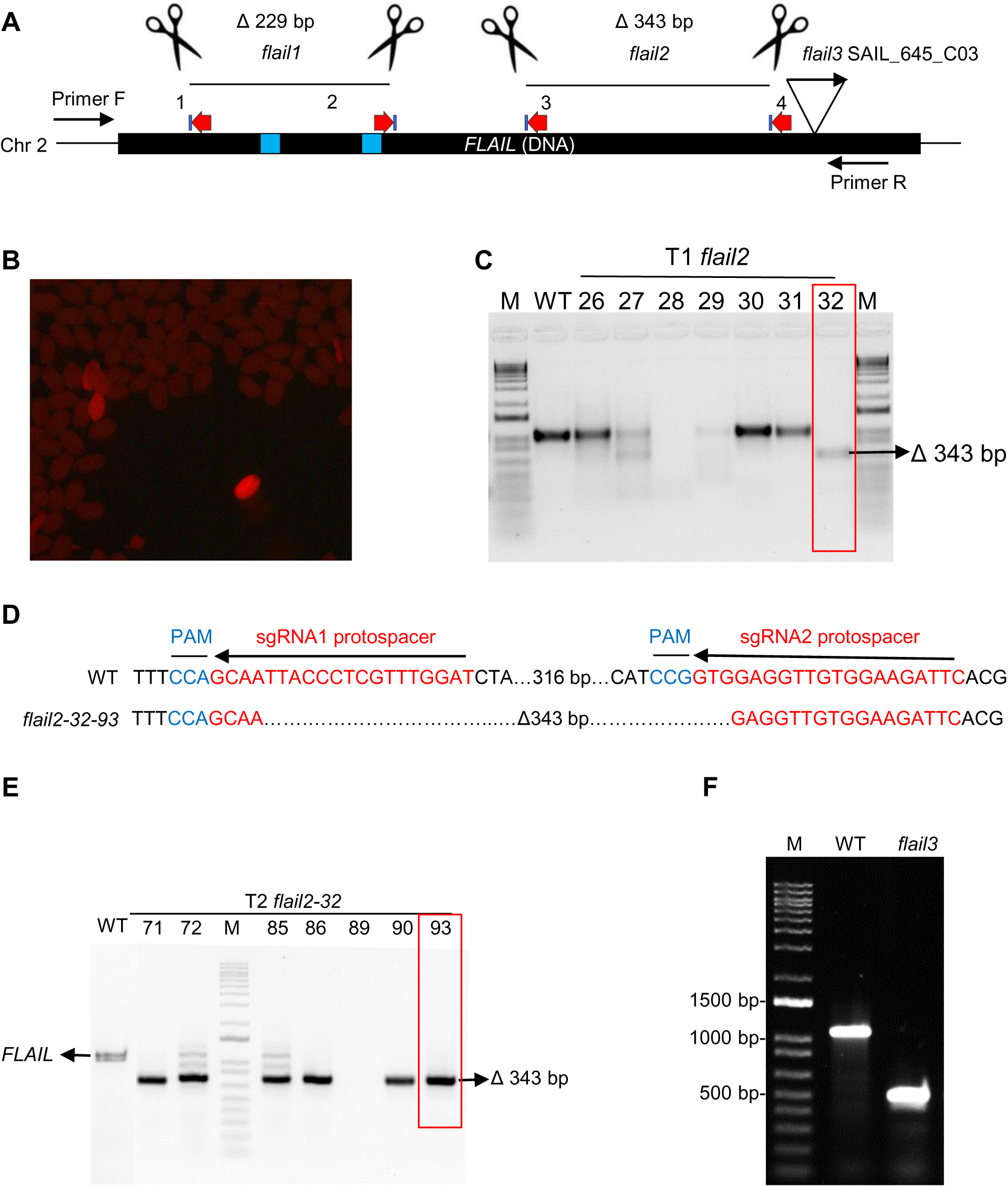
F*L*AIL dual-sgRNA approach and T-DNA insertion genotyping. **A** Schematic representation of targeted gene *FLAIL* with locations of two Cas9-directed mutants *flail1*, *flail2*, and one T-DNA mutant *flail3*. Locations of dual-sgRNA target sites were shown in red arrow, primer pair F/R (Mlo2478/2479) were used for PCR testing deletion in *flail1* and *flail2*, triangle indicated the putative position of T-DNA insertion at the *FLAIL* locus. Blue boxes indicated two sORFs. **B** Similar to the previous generation of *flail1*, *FLAIL2* construct containing pKIR1.1 enabled red fluorescence selection of *flail2* seeds from T1 plants after transformation by floral dipping. **C** Genotyping of individual T1 plant with red fluorescence, Leaf genomic DNA of T1 plants were PCR amplified. Expected size of PCR product for deletion was 343 bp in *flail2* line. *flail2-32* in red box was selected for next generation. **D** In T2 selection, the null mutant *flail2-32-93* without red fluorescence in seeds represented Cas9-free plants, indicated by red box in **E**. PAM was shown in blue, sgRNA protospacers were in red, and deleted bases were replaced by dots. **E** PCR analysis of T2 lines. The expected size of the wild-type *FLAIL* amplicon and the deletion size between Cas9 cut sites in *flail2-32-93* were indicated. This inherited Cas9-null segregation line was used for data analysis in Fig. 1. **F** Genotyping of the T-DNA insertion mutant *flail3*. Gel sample order (from left to right): lane 1, marker; lane 2, WT; lane 3, *flail3*, with primer set, Mlo1788 + Mlo1789 + Mlo37; Results clearly showed that *flail3* line was homozygous since WT was the only line giving WT band and *flail3* line gave the only T-DNA insertion band.

**Fig. S5.**
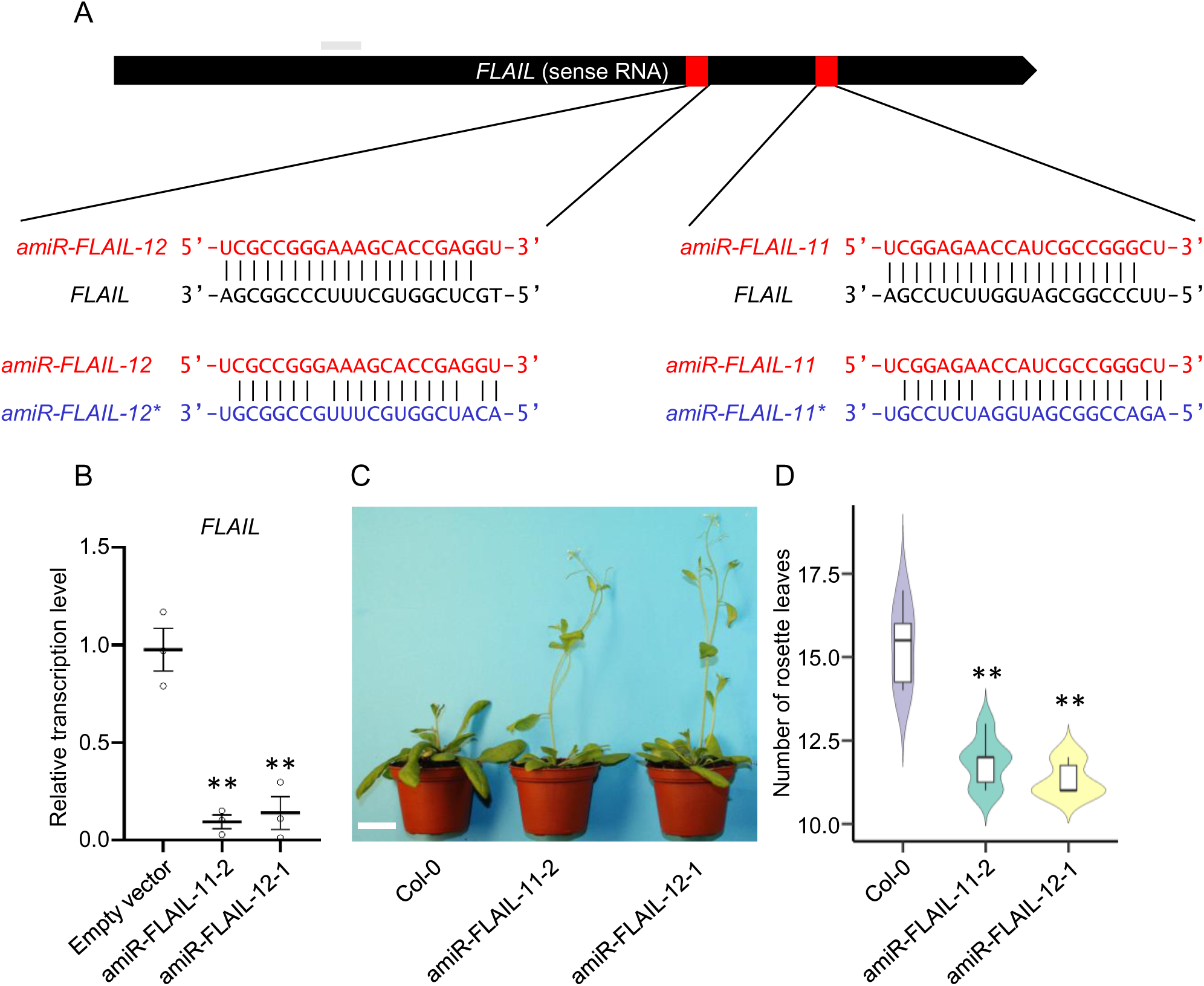
F*L*AIL lncRNA functions as a flowering repressor. **A** Sequences and structures of amiRNA duplexes and the target sites of *amiR-FLAILs* and *amiR-FLAIL*s*. Upper panel, schematic representation of transcribed sense *FLAIL* RNA. Lower panel, sequences of amiRNA duplexes, amiRNA (in red) target sites, and potential amiRNA* (in blue) on cognate sense *FLAIL* (in black). **B** Gene expression of *sense FLAIL* in *amiR-FLAIL* plants by RT-qPCR. Transcript levels were normalized to *UBQ* expression levels. Two representative lines for amiRNA designed transgenic plants were selected. Y-axis showed relative values compared to the expression level of empty vector transformed plants. Grey bars depicted the relative positions of primers used for RT-qPCR analyses. Transcript levels were normalized to *UBQ* expression levels. Error bars represented s.e.m (n = 3 independent 14-day seedling pools). **, *p* value < 0.01 analyzed by Student’s t-test. **C** Morphological phenotypes of 4-week-old plants of Col-0 and two *amiR-FLAIL* plants under a 16-h light/8-h dark growth condition. Scale bar: 2 cm. **D** Violin graph showed number of rosette leaves after appearance of the first flower bud in Col-0. Data represented the mean of six independent experiments. Boxes spanned the first to third quartile, bold black lines indicated median value for each group and whiskers represented the minimum and maximum values. **, *p* value < 0.01 was indicated by Student’s t-test compared to Col-0.

**Fig. S6.**
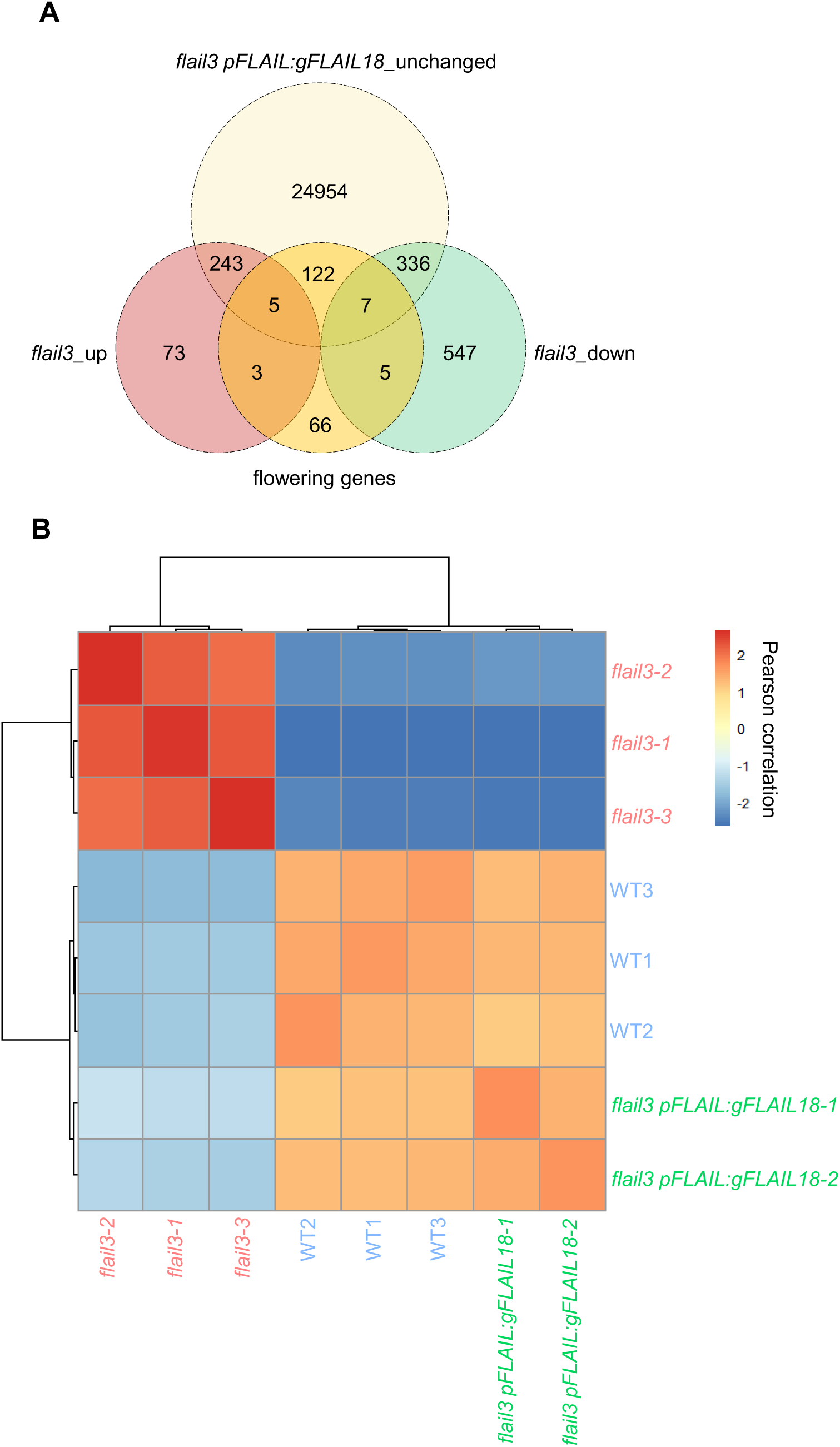
Genome-wide effects of *FLAIL* on gene expression by RNA-seq. **A** Venn diagram of flowering genes overlapped with upregulated and downregulated genes in the *flail3* mutant and with unchanged genes in the *flail3 pFLAIL:gFLAIL18* complementation plant. **B** Reproducibility of all genes shown in Fig. 3 from RNA-seq data was demonstrated by clustered heatmap of Pearson correlation coefficients over all independent replicates of RNA-seq in WT, *flail3* mutant, and *flail3 pFLAIL gFLAIL18* plants. Darker red denoted higher correlation and darker blue represented low reproducibility.

**Fig. S7.**
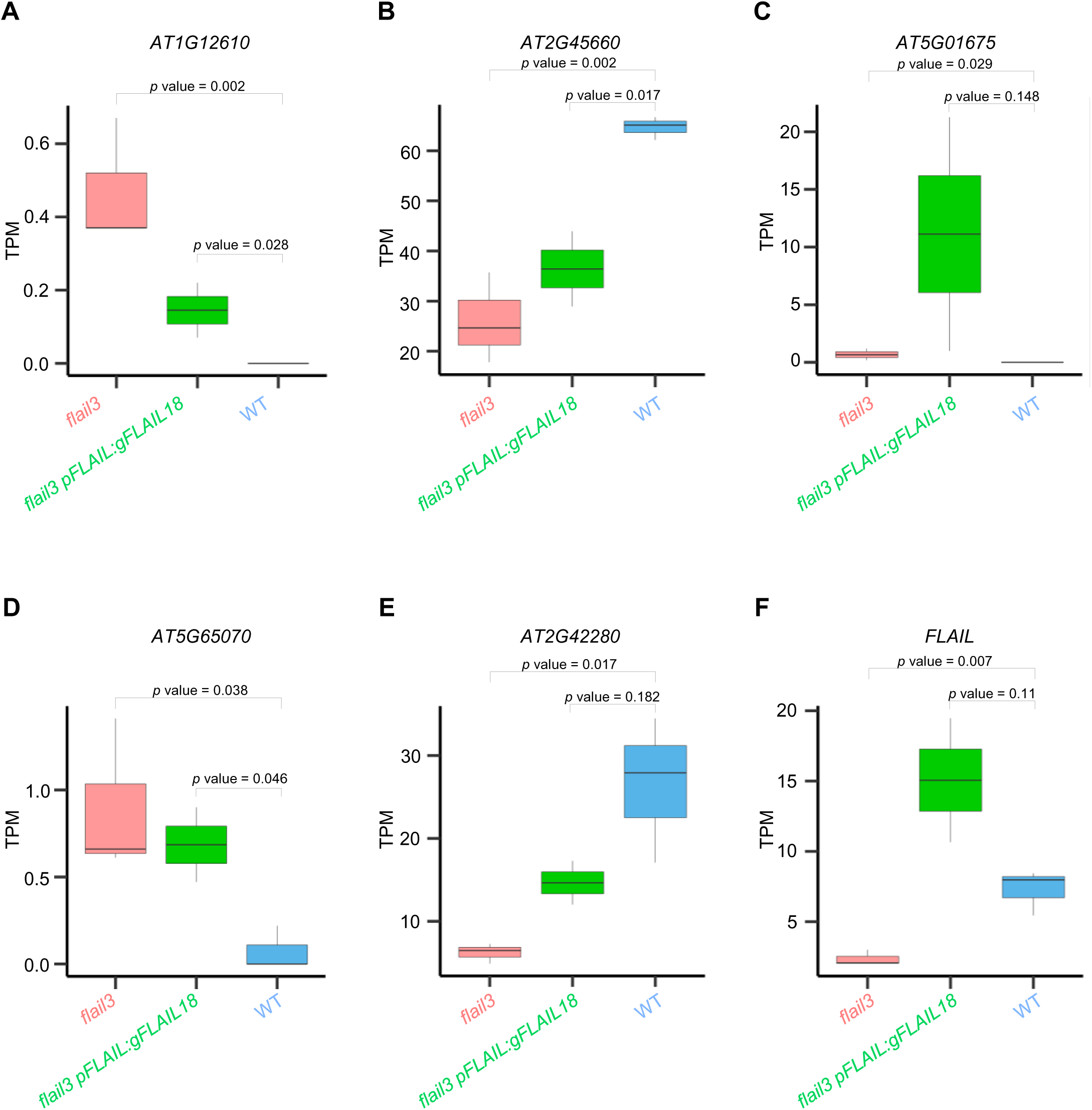
F*L*AIL regulates flowering related genes in *trans.* **A-F** Genome browser screenshots illustrating the expression of dysregulated flowering genes in *flail3* were most fully rescued in complementation line. Normalized read counts (TPM from RNA-seq) were used for differentially expressed flowering genes in WT, *flail3*, and *flail3 pFLAIL:gFLAIL18* plants. Boxes spanned the first to third quartile, bold black lines indicated median value for each group and whiskers represented the minimum and maximum values. *p* value was denoted by Students’ t-test.

**Fig. S8.**
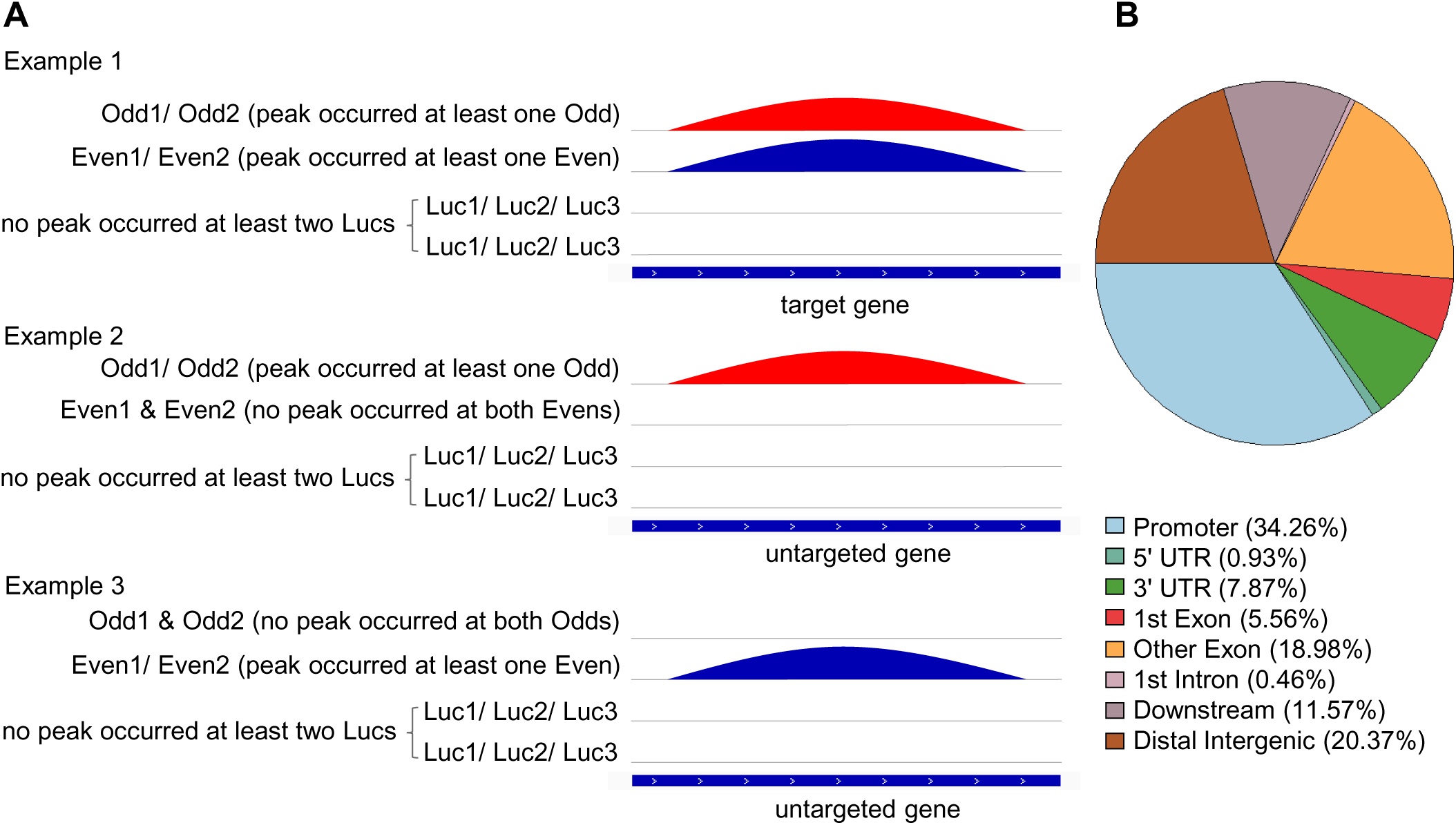
Strategy to identify genome-wide binding profile of *FLAIL* analyzed by ChIRP-seq. **A** Illustration of *FLAIL* peaks called by CCAT3.0. Only peaks that occurred at target genes from both Even and Odd probed DNA, but not from two or more independent experiments of Luc pools were considered significant enrichment (Example 1), peaks that only occurred from either Even or Odd pools were not considered *FLAIL* targets (Example 2 & 3). **B** A pie chart that was generated by ChIPseeker represented distribution of *FLAIL* bound regions in the genome. A total of 210 *FLAIL* bound regions were annotated according to the genomic distribution and *FLAIL* was enriched predominantly in promoter regions.

**Fig. S9.**
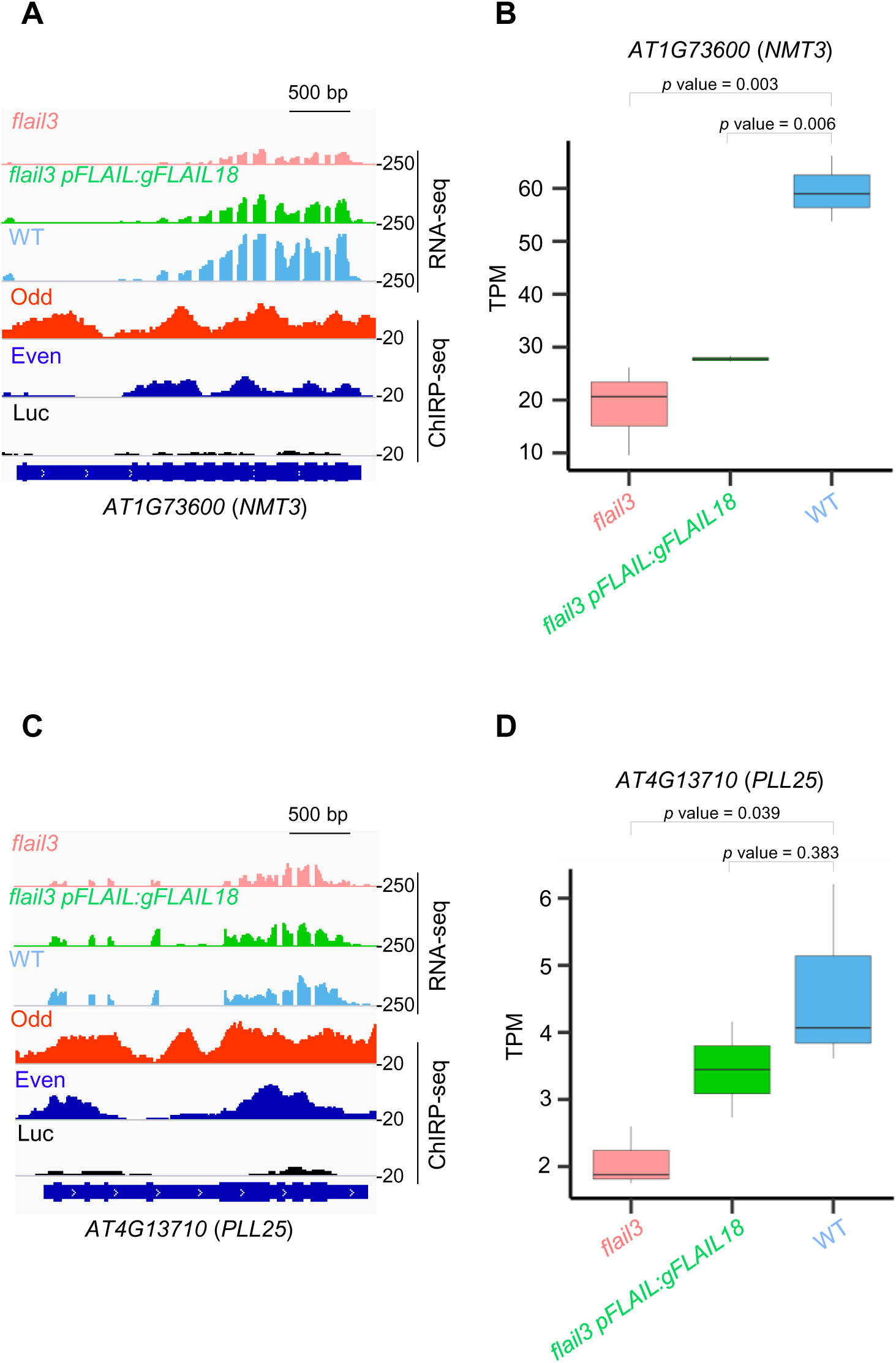
F*L*AIL binds chromatin regions of flowering genes for gene regulation. **A, C** Genome browser screenshots illustrating two more *FLAIL* bound flowering targets *PLL25* and *NMT3* in RNA-seq (lane 1-3) and ChIRP-seq (lane 4-6). Their expression that was downregulated in *flail3* (lane 1) can be partially or fully rescued in complementation line (lane 2). Both Odd and Even probes identified chromatin binding regions of *FLAIL* in two more flowering genes *NMT3* and *PLL25* compared to *Luc* probes. *FLAIL* binding peaks were called by CCAT3.0. **B, D** Normalized read counts (TPM from RNA-seq) for differentially expressed (DE) flowering genes in WT, *flail3*, and *flail3 pFLAIL:gFLAIL18* plants (bottom panel). Boxes spanned the first to third quartile, bold black lines indicated median value for each group and whiskers represented the minimum and maximum values. *p* value was denoted by Students’ t-test.

**Fig. S10.**
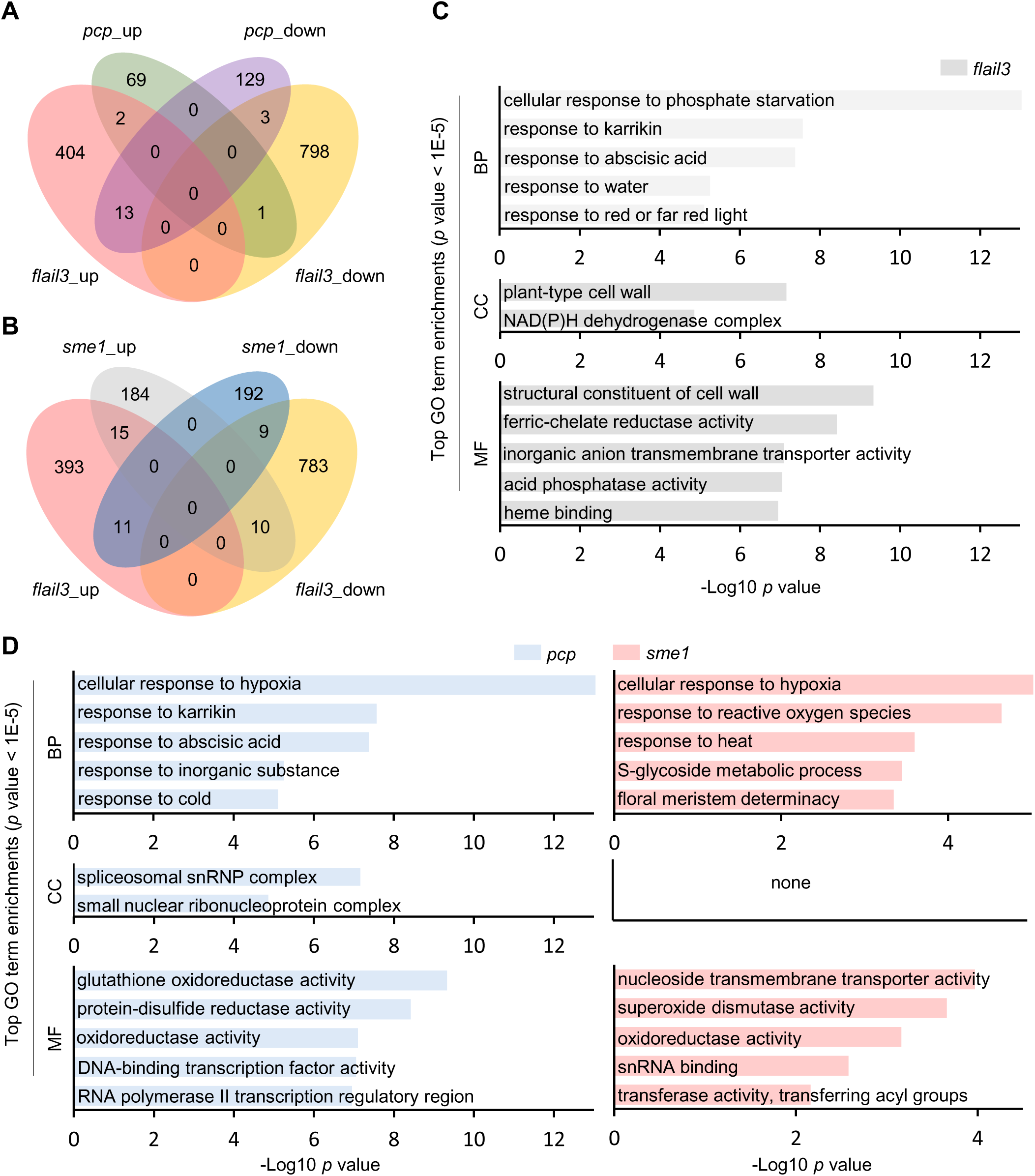
F*L*AIL regulates flowering independent of *PCP.* **A, B** Venn diagram showing overlapping differentially expressed genes (up/down) sets between *flail3* and *pcp* (also called *sme1*) mutants. **C, D** GO analysis of the DEGs in RNA-seq data of *flail3* and *pcp /sme1* mutants. Y-axis indicated the GO categories including biological process (BP), cellular component (CC) and molecular function (MF); X-axis showed -Log10 *p* value with the cutoff 0.05. Highly enriched GO terms of dysregulated mRNAs analyzed by the Metascape with 5 top enrichment scores.

